# Alpha-ketoamides as broad-spectrum inhibitors of coronavirus and enterovirus replication

**DOI:** 10.1101/2020.02.10.936898

**Authors:** Linlin Zhang, Daizong Lin, Yuri Kusov, Yong Nian, Qingjun Ma, Jiang Wang, Albrecht von Brunn, Pieter Leyssen, Kristina Lanko, Johan Neyts, Adriaan de Wilde, Eric J. Snijder, Hong Liu, Rolf Hilgenfeld

**Affiliations:** Institute of Biochemistry, Center for Structural and Cell Biology in Medicine, University of Lübeck, 23562 Lübeck, Germany; German Center for Infection Research (DZIF), Hamburg - Lübeck - Borstel - Riems Site, University of Lübeck, Lübeck, Germany; Shanghai Institute of Materia Medica, 201203 Shanghai, China; Max von Pettenkofer Institute, Ludwig-Maximilians-University Munich, 80336 Munich, Germany; Rega Institute for Medical Research, University of Leuven, 3000 Leuven, Belgium; Leiden University Medical Center, 2333 ZA Leiden, The Netherlands

**Author notes:** Corresponding Author Prof. Rolf Hilgenfeld, Institute of Biochemistry, University of Lübeck, Ratzeburger Allee 160, 23562 Lübeck Germany; Prof. Hong Liu, Shanghai Institute of Materia Medica, Zu Chong Zhi Road 555, Shanghai, 201203 China. These authors contributed equally (Linlin Zhang, Daizong Lin).

**Keywords:** coronavirus main protease, enterovirus 3C protease, antiviral drug design, X-ray crystallography, SARS, MERS, human coronavirus 229E, human coronavirus NL63, Wuhan pneumonia coronavirus, 2019 novel coronavirus, enterovirus A71, Coxsackievirus B3, enterovirus D68, human rhinovirus

## Abstract

The main protease of coronaviruses and the 3C protease of enteroviruses share a similar active-site architecture and a unique requirement for glutamine in the P1 position of the substrate. Because of their unique specificity and essential role in viral polyprotein processing, these proteases are suitable targets for the development of antiviral drugs. In order to obtain near-equipotent, broad-spectrum antivirals against alphacoronaviruses, betacoronaviruses, and enteroviruses, we pursued structure-based design of peptidomimetic α-ketoamides as inhibitors of main and 3C proteases. Six crystal structures of protease:inhibitor complexes were determined as part of this study. Compounds synthesized were tested against the recombinant proteases as well as in viral replicons and virus-infected cell cultures; most of them were not cell-toxic. Optimization of the P2 substituent of the α-ketoamides proved crucial for achieving near-equipotency against the three virus genera. The best near-equipotent inhibitors, **11u** (P2 = cyclopentylmethyl) and **11r** (P2 = cyclohexylmethyl), display low-micromolar EC_50_ values against enteroviruses, alphacoronaviruses, and betacoronaviruses in cell cultures. In Huh7 cells, **11r** exhibits three-digit picomolar activity against Middle East Respiratory Syndrome coronavirus.

## INTRODUCTION

Seventeen years have passed since the outbreak of severe acute respiratory syndrome (SARS) in 2003, but there is yet no approved treatment for infections with the SARS coronavirus (SARS-CoV).^1^ One of the reasons is that despite the devastating consequences of SARS for the affected patients, the development of an antiviral drug against this virus would not be commercially viable in view of the fact that the virus has been rapidly contained and did not reappear since 2004. As a result, we were empty-handed when the Middle-East respiratory syndrome coronavirus (MERS-CoV), a close relative of SARS-CoV, emerged in 2012.^2^ MERS is characterized by severe respiratory disease, quite similar to SARS, but in addition frequently causes renal failure^3^. Although the number of registered MERS cases is low (2494 as of November 30, 2019; www.who.int), the threat MERS-CoV poses to global public health may be even more serious than that presented by SARS-CoV. This is related to the high case-fatality rate (about 35%, compared to 10% for SARS), and to the fact that MERS cases are still accumulating seven years after the discovery of the virus, whereas the SARS outbreak was essentially contained within 6 months. The potential for human-to-human transmission of MERS-CoV has been impressively demonstrated by the 2015 outbreak in South Korea, where 186 cases could be traced back to a single infected traveller returning from the Middle East.^4^ SARS-like coronaviruses are still circulating in bats in China,^5-8^ from where they may spill over into the human population; this is probably what caused the current outbreak of atypical pneumonia in Wuhan, which is linked to a seafood and animal market. The RNA genome (GenBank accession code: MN908947.2; http://virological.org/t/initial-genome-release-of-novel-coronavirus/319, last accessed on January 11, 2020) of the new betacoronavirus features around 82% identity to that of SARS-CoV.

In spite of the considerable threat posed by SARS-CoV and related viruses, as well as by MERS-CoV, it is obvious that the number of cases so far does not warrant the commercial development of an antiviral drug targeting MERS- and SARS-CoV even if a projected steady growth of the number of MERS cases is taken into account. A possible solution to the problem could be the development of broad-spectrum antiviral drugs that are directed against the major viral protease, a target that is shared by all coronavirus genera as well as, in a related form, by members of the large genus *Enterovirus* in the picornavirus family. Among the members of the genus *Alphacoronavirus* are the human coronaviruses (HCoV) NL63 (ref. 9) and 229E^10^ that usually cause only mild respiratory symptoms in otherwise healthy individuals, but are much more widespread than SARS-CoV or MERS-CoV. Therapeutic intervention against alphacoronaviruses is indicated in cases of accompanying disease such as cystic fibrosis^11^ or leukemia,^12^ or certain other underlying medical conditions.^13^ The enteroviruses include pathogens such as EV-D68, the causative agent of the 2014 outbreak of the “summer flu” in the US,^14^ EV-A71 and Coxsackievirus A16 (CVA16), the etiological agents of Hand, Foot, and Mouth Disease (HFMD),^15^ Coxsackievirus B3 (CVB3), which can cause myocardic inflammation,^16^ and human rhinoviruses (HRV), notoriously known to lead to the common cold but also capable of causing exacerbations of asthma and COPD.^17^ Infection with some of these viruses can lead to serious outcome; thus, EV-D68 can cause polio-like disease,^18^ and EV-A71 infection can proceed to aseptic meningitis, encephalitis, pulmonary edema, viral myocarditis, and acute flaccid paralysis.^15,19-20^ Enteroviruses cause clinical disease much more frequently than coronaviruses, so that an antiviral drug targeting both virus families should be commercially viable.

However, enteroviruses are very different from coronaviruses. While both of them have a single-stranded RNA genome of positive polarity, that of enteroviruses is very small (just 7 - 9 kb) whereas coronaviruses feature the largest RNA genome known to date (27 - 34 kb). Enteroviruses are small, naked particles, whereas coronaviruses are much larger and enveloped. Nevertheless, a related feature shared by these two groups of viruses is their type of major protease,^21^ which in the enteroviruses is encoded by the 3C region of the genome (hence the protease is designated 3C^pro^). In coronaviruses, non-structural protein 5 (Nsp5) is the main protease (M^pro^). Similar to the enteroviral 3C^pro^, it is a cysteine protease in the vast majority of cases and has therefore also been called “3C-like protease” (3CL^pro^). The first crystal structure of a CoV M^pro^ or 3CL^pro^ (ref. 22) revealed that two of the three domains of the enzyme together resemble the chymotrypsin-like fold of the enteroviral 3C^pro^, but there is an additional α-helical domain that is involved in the dimerization of the protease (Fig. 1A). This dimerization is essential for the catalytic activity of the CoV M^pro^, whereas the enteroviral 3C^pro^ (Fig. 1B) functions as a monomer. Further, the enteroviral 3C^pro^ features a classical Cys…His…Glu/Asp catalytic triad, whereas the CoV M^pro^ only has a Cys…His dyad.^22^ Yet, there are a number of common features shared between the two types of proteases, in particular their almost absolute requirement for Gln in the P1 position of the substrate and space for only small amino-acid residues such as Gly, Ala, or Ser in the P1’ position, encouraging us to explore the coronaviral M^pro^ and the enteroviral 3C^pro^ as a common target for the design of broad-spectrum antiviral compounds. The fact that there is no known human protease with a specificity for Gln at the cleavage site of the substrate increases the attractivity of this viral target, as there is hope that the inhibitors to be developed will not show toxicity versus the host cell. Indeed, neither the enterovirus 3C^pro^ inhibitor rupintrivir, which was developed as a treatment of the common cold caused by HRV, nor the peptide aldehyde inhibitor of the coronavirus M^pro^ that was recently demonstrated to lead to complete recovery of cats from the normally fatal infection with Feline Infectious Peritonitis Virus (FIPV), showed any toxic effects on humans or cats, respectively.^23,24^

**Figure 1:**
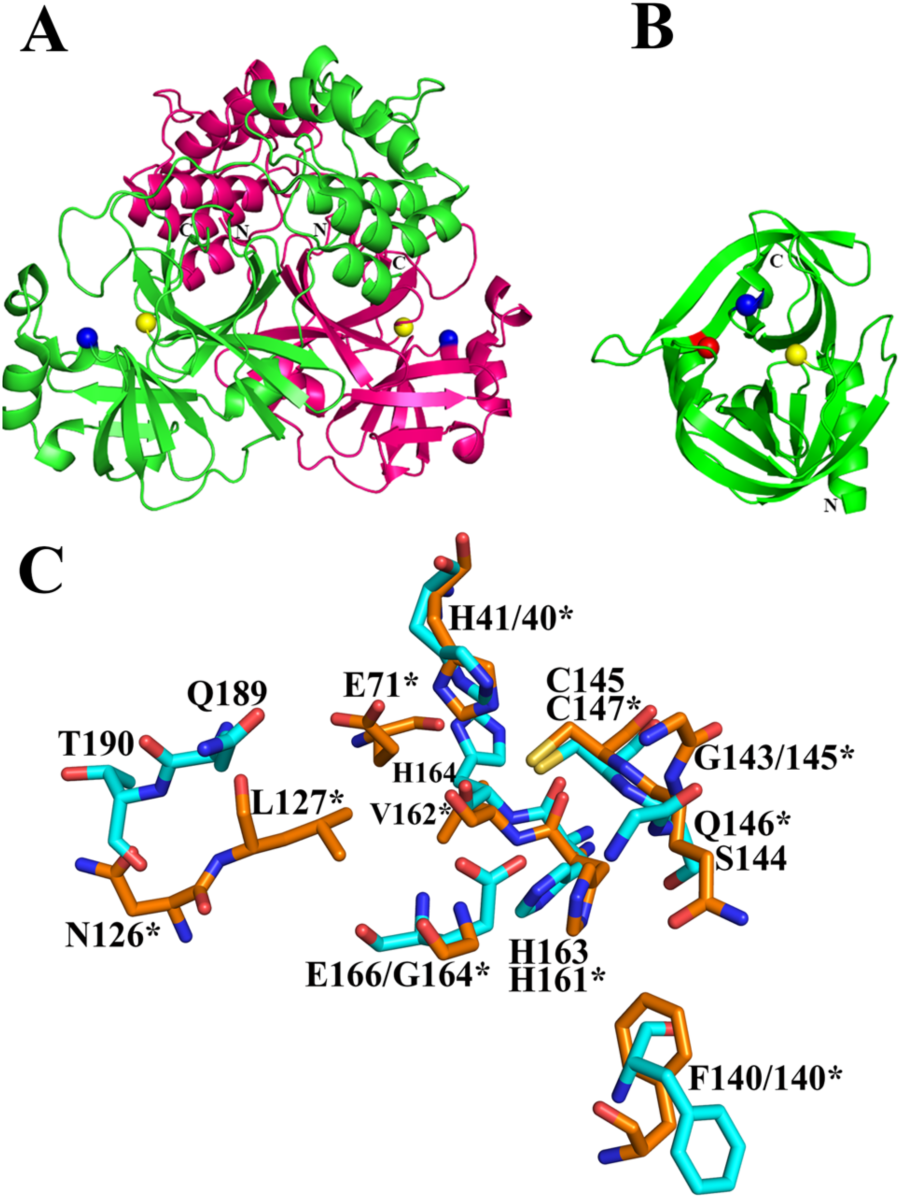
Crystal structures of SARS-CoV main protease (M^pro^, ref. 26; PDB entry 2BX4) and Coxsackivirus B3 3C protease (3C^pro^; Tan et al., unpublished; PDB entry 3ZYD). Catalytic residues are indicated by spheres (yellow, Cys; blue, His; red: Glu). A. The coronavirus M^pro^ is a homodimer, with each monomer comprising three domains. B. The structure of the monomeric CVB3 3C^pro^ resembles the N-terminal two domains of the SARS-CoV M^pro^. Structure is on the same scale as image A. C. Superimpostion of residues from the two structures involved in ligand binding. Superimposition was carried out by aligning the catalytic Cys-His pair of each protease. Residues of the SARS-CoV M^pro^ are shown with carbon atoms in cyan, CVB3 3C^pro^ residues have orange carbons and are labeled with an asterisk (*).

We chose the chemical class of peptidomimetic α-ketoamides to assess the feasibility of achieving antiviral drugs targeting coronaviruses and enteroviruses with near-equipotency. Here we describe the structure-based design, synthesis, and evaluation of inhibitory activity of a series of compounds with broad-spectrum activities afforded by studying the structure-activity relationships mainly with respect to the P2 position of the peptidomimetics. One of the compounds designed and synthesized exhibits excellent activity against MERS-CoV.

## RESULTS

### Structure-based design of α-ketoamides

Our efforts to design novel α-ketoamides as broad-spectrum inhibitors of coronavirus M^pro^s and enterovirus 3C^pro^s started with a detailed analysis of the following crystal structures of unliganded target enzymes: SARS-CoV M^pro^ (ref. 25-27; PDB entries 1UJ1, 2BX3, 2BX4); bat coronavirus HKU4 M^pro^ as a surrogate for the closely related MERS-CoV protease (our unpublished work (Ma, Xiao et al.; PDB entry 2YNA; see also ref. 27); HCoV-229E M^pro^ (ref. 27,28; PDB entry: 1P9S); Coxsackievirus B3 3C^pro^ (our unpublished work; Tan et al., PDB entry 3ZYD); enterovirus D68 3C^pro^ (ref. 29; PDB entry: 3ZV8); and enterovirus A71 3C^pro^ (ref. 30; PDB entry: 3SJK). During the course of the present study, we determined crystal structures of a number of lead α-ketoamide compounds in complex with SARS-CoV M^pro^, HCoV-NL63 M^pro^, and CVB3 3C^pro^, in support of the design of improvements in the next round of lead optimization. Notably, unexpected differences between alpha- and betacoronavirus M^pro^ were found in this study. The structural foundation of these was elucidated in detail in a subproject involving the M^pro^ of HCoV NL63; because of its volume, this work will be published separately (Zhang et al., in preparation) and only some selected findings are referred to here. The main protease of the newly discovered coronavirus linked to the Wuhan outbreak of respiratory illness is 96% identical (98% similar) in amino-acid sequence to that of SARS-CoV M^pro^ (derived from the RNA genome of BetaCoV/Wuhan/IVDC-HB-01/2019, Genbank accession code: MN908947.2; http://virological.org/t/initial-genome-release-of-novel-coronavirus/319, last accessed on January 11, 2020), so all results reported here for inhibition of SARS-CoV will most likely also apply to the new virus.

As the proteases targeted in our study all specifically cleave the peptide bond following a P1-glutamine residue (HCoV-NL63 M^pro^ uniquely also accepts P1 = His at the Nsp13/Nsp14 cleavage site^31^), we decided to use a 5-membered ring (γ-lactam) derivative of glutamine (henceforth called GlnLactam) as the P1 residue in all our α-ketoamides (see Scheme 1). This moiety has been found to be a good mimic of glutamine and enhance the power of the inhibitors by up to 10-fold, most probably because compared to the flexible glutamine side-chain, the more rigid lactam leads to a reduction of the loss of entropy upon binding to the target protease.^29,32^ Our synthetic efforts therefore aimed at optimizing the substituents at the P1’, P2, and P3 positions of the α-ketoamides.

### Synthesis of α-ketoamides

Synthesis (Scheme 1) started with the dianionic alkylation of *N*-Boc glutamic acid dimethyl ester with bromoacetonitrile. As expected, this alkylation occurred in a highly stereoselective manner, giving **1** as the exclusive product. In the following step, the cyano group of **1** was subjected to hydrogenation. The *in-situ* cyclization of the resulting intermediate afforded the lactam **2**. The lactam derivative **3** was generated by removal of the protecting group of **2**. On the other hand, the amidation of acyl chloride and α-amino acid methyl ester afforded the intermediates **4**, which gave rise to the acids **5** via alkaline hydrolysis. The key intermediates **6** were obtained *via* the condensation of the lactam derivative **3** and the N-capped amino acids **5**. The ester group of compounds **6** was then reduced to the corresponding alcohol. Oxidation of the alcohol products **7** by Dess-Martin periodinane generated the aldehydes **8**, which followed by nucleophilic addition with isocyanides gave rise to compounds **9** under acidic conditions. Then, the α-hydroxyamides **10** were prepared by removing the acetyl group of compounds **9**. In the final step, the oxidation of the exposed alcohol group in compounds **10** generated our target α-ketoamides **11**.

The inhibitory potencies of candidate α-ketoamides were evaluated against purified recombinant SARS-CoV M^pro^, HCoV-NL63 M^pro^, CVB3 3C^pro^, and EV-A71 3C^pro^. The most potent compounds were further tested against viral replicons and against SARS-CoV, MERS-CoV, or a whole range of enteroviruses in cell culture-based assays (Tables 1 - 3 and Supplementary Table 1).

**Table 1:** Inhibition of viral proteases by α-ketoamides (IC_50_, μM)

| Compound No. | Formula | EV-A71 3C<sup>pro</sup> | CVB3 3C<sup>pro</sup> | SARS-CoV M<sup>pro</sup> | HCoV-NL63 M<sup>pro</sup> |
| --- | --- | --- | --- | --- | --- |
| 11a |  | 1.22 ± 0.12 | 6.56 ± 3.10 | 1.95 ± 0.24 | >50 |
| 11f |  | >50 | >50 | >50 | >50 |
| 11m |  | 2.32 ± 1.21 | 8.69 ± 1.28 | >50 | >50 |
| 11n |  | 13.80 ± 4.17 | 3.82 ± 1.21 | 0.33 ± 0.04 | 1.08 ± 0.09 |
| 11o |  | 3.24 ± 1.69 | 5.20 ± 0.28 | 8.50 ± 3.71 | >50 |
| 11p |  | 8.50 ± 1.81 | 8.93 ± 2.39 | 10.68 ± 7.34 | 17.37 ± 3.97 |
| 11q |  | >30 | 15.38 ± 4.12 | 6.27 ± 2.87 | 11.50 ± 5.05 |
| 11r |  | 1.69 ± 0.47 | 0.95 ± 0.15 | 0.71 ± 0.36 | 12.27 ± 3.56 |
| 11s |  | 18.47 ± 4.25 | 4.25 ± 1.40 | 0.24 ± 0.08 | 1.37 ± 0.35 |
| 11t |  | 10.81 ± 2.62 | 4.78 ± 1.13 | 1.44 ± 0.40 | 3.43 ± 2.45 |
| 11u |  | 4.73 ± 0.94 | 1.93 ± 0.43 | 1.27 ± 0.34 | 5.41 ± 2.31 |
| nd – not done |  |  |  |  |  |

**Table 2:** α-ketoamide-induced inhibition of subgenomic RNA synthesis using replicons in a cell-based assay (EC_50_, μM)

| Compound No. | EV-A71 | CVB3 | SARS-CoV |
| --- | --- | --- | --- |
| **11a** | $4.5 \pm 0.5$ | $4.5 \pm 0.5$ | $2.0 \pm 0.2$ |
| **11f** | $>>10$ | $>>10$ | $>40$ |
| **11m** | $>20$ | nd | nd |
| **11n** | $>>10$ | $>>10$ | $7.2 \pm 0.2$ |
| **11o** | $>20$ | nd | nd |
| **11p** | nd | nd | $>20$ |
| **11r** | $6.95 \pm 0.05$<br>($0.85 \pm 0.05$)* | $2.35 \pm 0.05$<br>($0.45 \pm 0.05$)* | $1.4 \pm 0.1$ |
| **11s** | $>>20$ | $18 \pm 2$ | $1.9 \pm 0.1$ |
| **11t** | $>20$<br>($10.9 \pm 0.1$)* | $4.3 \pm 0.2$<br>($2.2 \pm 0.2$)* | $6.7 \pm 0.2$ |
| **11u** | $8.9 \pm 0.1$<br>($3.65 \pm 0.15$)* | $5.1 \pm 0.1$<br>($4.9 \pm 2.6$)* | $3.6 \pm 0.1$ |

nd - not done; \* - values in brackets obtained by RNA-launched transfection.

**Table 3:** Cytotoxicity and antiviral activity of α-ketoamides against selected entero- and coronaviruses in a live-virus cell-based assay.

| Cells | Huh-T7 | Huh-T7 | Huh7 | Huh7 | Vero | Vero E6 | RD | RD | Hela Rh | Hela Rh | Hela Rh | Hela Rh |
| --- | --- | --- | --- | --- | --- | --- | --- | --- | --- | --- | --- | --- |
| Virus<br>(Strain) | -- | CVB3<br>(Nancy) | HCoV-229E | MERS-CoV | MERS-CoV | SARS-CoV | -- | EV-A71 | -- | EV-D68 | HRV2 | HRV14 |
| Compound | CC<sub>50</sub> (μM) | EC<sub>50</sub> (μM) | EC<sub>50</sub> (μM) | EC<sub>50</sub> (μM) | EC<sub>50</sub> (μM) | EC<sub>50</sub> (μM) | CC<sub>50</sub> (μM) | EC<sub>50</sub> (μM) | CC<sub>50</sub> (μM) | EC<sub>50</sub> (μM) | EC<sub>50</sub> (μM) | EC<sub>50</sub> (μM) |
| 11a | >265 | 14 ± 1 | 11.8 ± 2.1 | 0.0048 ± 0.001 | 13.4 | 5.8 | 97 ± 7 | 9.8 ± 1.1 | 35.5 ± 2.7 | 0.48 ± 0.09 | 5.6 ± 0.7 | 4.3 ± 0.1 |
| 11n | >188 | 24 ± 5 | 0.6 ± 0.1 | 0.0046 ± 0.001 | 9.2 | 14.2 | 115 ± 14 | 72 ± 14 | 103 ± 5 | 4.44 ± 0.02 | 8.9 ± 2.5 | 13.40 ± 0.01 |
| 11r | 43 ± 2 | 2.77 ± 0.03 | 1.8 ± 0.6 | 0.0004 ± 0.0003 | 5.0 ± 0.4 | 2.1 ± 1.2 | 17.3 ± 2.9 | 3.7 ± 0.2 | 10.9 ± 1.8 | 0.48 ± 0.09 | 0.64 ± 0.05 | 0.70 ± 0.09 |
| 11s | >189 | 10.4 ± 0.1 | 1.3 ± 0.4 | 0.08 ± 0.01 | 10.9 ± 1.7 | 18.4 ± 6.7 | 106 ± 8 | 46 ± 7 | 103 ± 2 | nd | 3.8 ± 0.2 | 5.5 ± 1.2 |
| 11t | 124 ± 1 | 10.4 ± 0.1 | nd | 0.02 ± 0.01 | 9.8 ± 1.5 | 7.0 ± 2.3 | 100 ± 2 | 41 ± 3 | 65 ± 7 | nd | 3.7 ± 0.4 | 4.2 ± 0.1 |
| 11u | 47.5 ± 2.3 | 8.1 ± 0.8 | 2.5 ± 0.6 | 0.007 ± 0.006 | 11.1 ± 0.2 | 4.9 ± 1.2 | 30.8 ± 0.1 | 4.20 ± 0.02 | 33.2 ± 0.9 | nd | 0.51 ± 0.05 | 0.42 ± 0.02 |
| nd – not done |  |  |  |  |  |  |  |  |  |  |  |  |

**Scheme 1:**
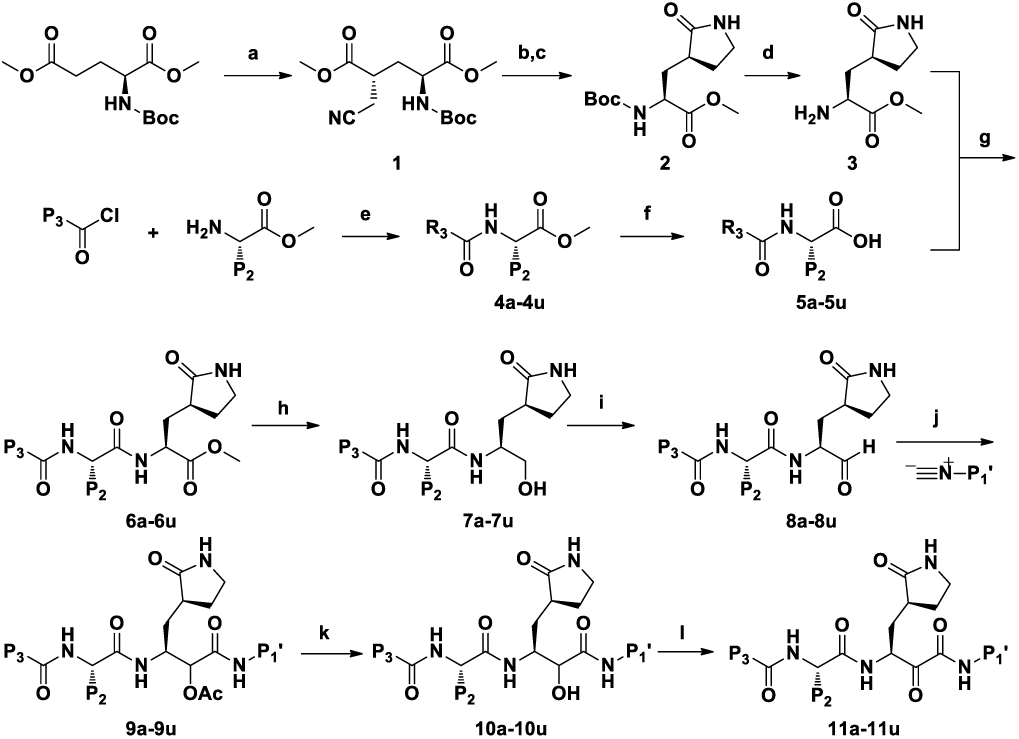
Synthesis of α-ketoamides. Reaction conditions: (a) BrCH_2_CN, LiHMDS, THF; (b) PtO_2_, H_2_, CHCl_3_, MeOH; (c) NaOAc, MeOH; (d) TFA, CH_2_Cl_2_; (e) TEA, CH_2_Cl_2_; (f) 1 M NaOH or LiOH, MeOH; (g) HATU, DMF; (h) NaBH_4_, MeOH; (i) DMP, NaHCO_3_, CH_2_Cl_2_; (j) isocyanide, AcOH, CH_2_Cl_2_; (k) 1 M NaOH or LiOH, MeOH; (l) DMP, NaHCO_3_, CH_2_Cl

### Viral replicons

To enable the rapid and biosafe screening of antivirals against corona- and enteroviruses, a non-infectious, but replication-competent SARS-CoV replicon was used^33^ along with subgenomic replicons of CVB3^34^ and EV-A71 (kind gift of B. Zhang, Wuhan, China). The easily detectable reporter activity (firefly or Renilla luciferase) of these replicons has previously been shown to reflect viral RNA synthesis.^33-35^ *In-vitro* RNA transcripts of the enteroviral replicons were also used for transfection. For the SARS-CoV replicon containing the CMV promoter, only the plasmid DNA was used for transfection.

### Initial inhibitor design steps

The initial compound to be designed and synthesized was **11a**, which carries a cinnamoyl N-cap in the P3 position, a benzyl group in P2, the glutamine lactam (GlnLactam) in P1, and benzyl in P1’ (Table 1). This compound showed good to mediocre activities against recombinant SARS-CoV M^pro^ (IC_50_ = 1.95 μM; for all compounds, see Tables 1 - 3 for standard deviations), CVB3 3C^pro^ (IC_50_ = 6.6 μM), and EV-A71 3C^pro^ (IC_50_ = 1.2 μM), but was surprisingly completely inactive (IC_50_ > 50 μM) against HCoV-NL63 M^pro^. These values were mirrored in the SARS-CoV and in the enterovirus replicons (Table 2). In virus-infected cell cultures, the results obtained were also good to mediocre (Table 3): SARS-CoV (EC_50_ = 5.8 μM in Vero E6 cells), MERS-CoV (EC_50_ = 0.0047 μM in Huh7 cells), HCoV 229E (EC_50_ = 11.8 μM in Huh7 cells), or a host of enteroviruses (EC_50_ = 9.8 μM against EV-A71 in RD cells; EC_50_ = 0.48 μM against EV-D68 in HeLa Rh cells; EC_50_ = 5.6 μM against HRV2 in HeLa Rh cells). In all cell types tested, the compound generally proved to be non-toxic, with selectivity indices (CC_50_/EC_50_) usually >10 (Table 3).

Crystal structures of compound **11a** in complex with SARS-CoV M^pro^, HCoV-NL63 M^pro^, and CVB3 3C^pro^ demonstrated that the α-keto-carbon is covalently linked to the active-site Cys (no. 145, 144, and 147, resp.) of the protease (Fig. 2, 3a-c). The resulting thiohemiketal is in the *R* configuration in the SARS-CoV and HCoV-NL63 M^pro^ but in the *S* configuration in the CVB3 3C^pro^ complex. The reason for this difference is that the oxygen atom of the thiohemiketal accepts a hydrogen bond from the catalytic His40 in the CVB3 protease, rather than from the main-chain amides of the oxyanion hole as in the SARS-CoV and HCoV-NL63 enzymes (Fig. 3a,b,c insets). It is remarkable that we succeeded in obtaining a crystal structure of compound **11a** in complex with the HCoV-NL63 M^pro^, even though it has no inhibitory effect on the activity of the enzyme (IC_50_ > 50 μM), (Fig. 2c). Apparently, the compound is able to bind to this M^pro^ in the absence of peptide substrate, but cannot compete with substrate for the binding site due to low affinity. A similar observation has been made in one of our previous studies, where we were able to determine the crystal structure of a complex between the inactive Michael-acceptor compound **SG74** and the EV-D68 3C^pro^ (ref. 29; PDB entry 3ZV9).

**Figure 2:**
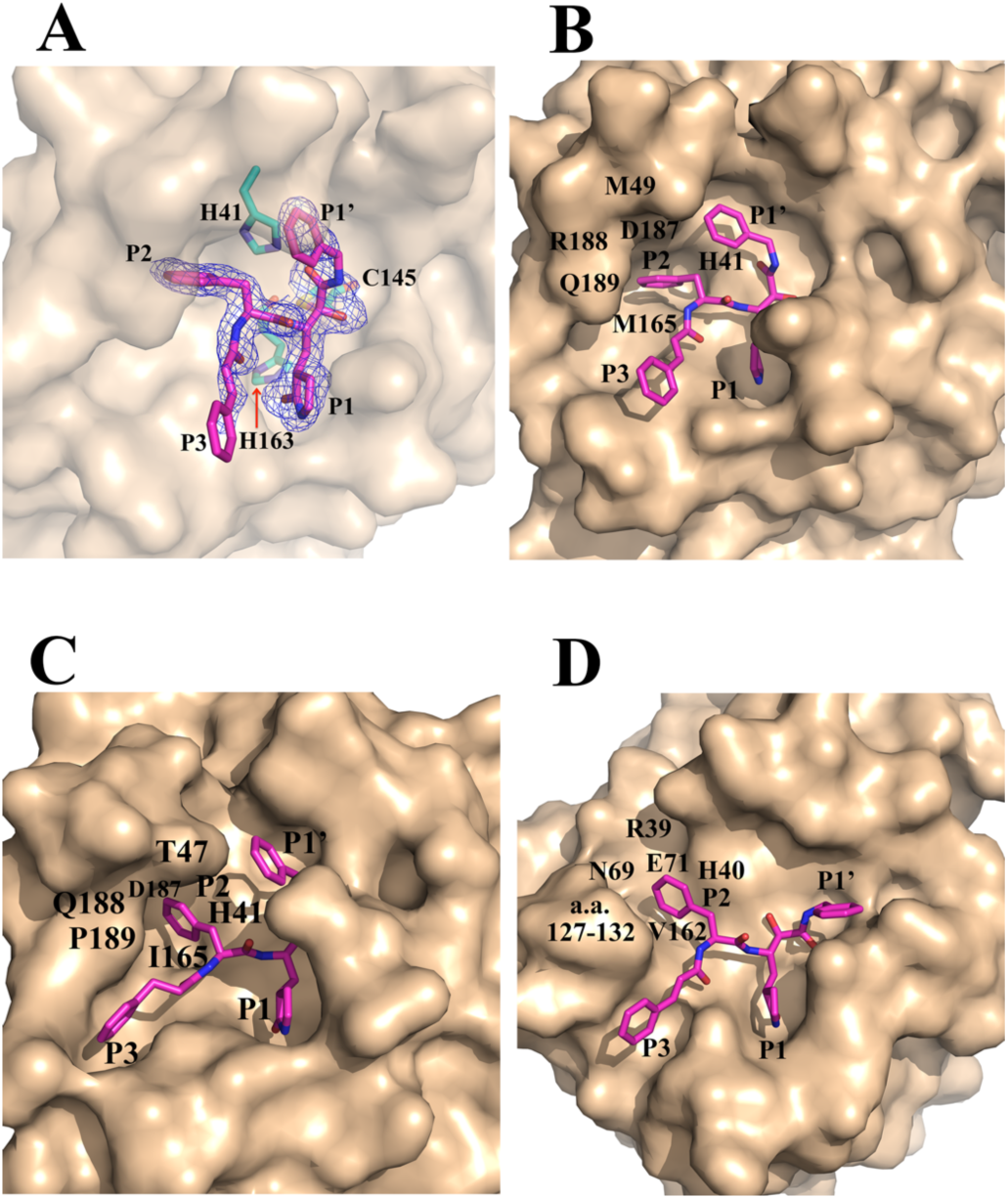
Fit of compound **11a** (pink carbon atoms) to the target proteases (wheat surfaces) as revealed by X-ray crystallography of the complexes. A. F_o_ - F_c_ difference density (contoured at 3σ) for **11a** in the substrate-binding site of the SARS-CoV M^pro^ (transparent surface). Selected side-chains of the protease are shown with green carbon atoms. B. Another view of **11a** in the substrate binding site of the SARS-CoV M^pro^. Note the “lid” formed by residue Met49 and its neighbors above the S2 pocket. C. **11a** in the substrate-binding site of HCoV-NL63 M^pro^. Because of the restricted size of the S2 pocket, the P2 benzyl group of the compound cannot enter deeply into this site. Note that the S2 pocket is also covered by a “lid” centred around Thr47. D. **11a** in the substrate-binding site of the CVB3 3C^pro^. The S2 site is large and not covered by a “lid”.

**Figure 3:**
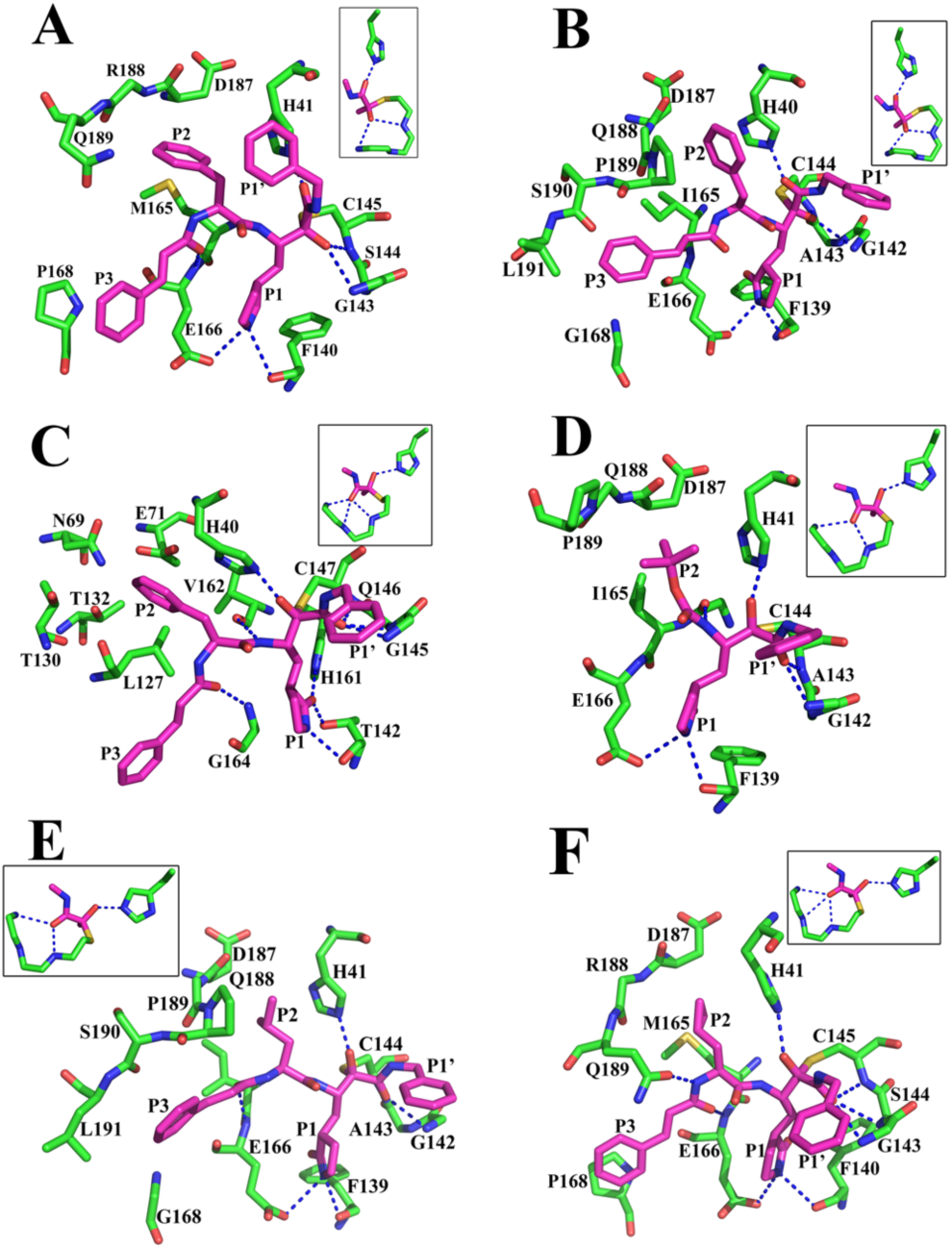
Detailed interactions of peptidomimetic α-ketoamides (pink carbon atoms) with target proteases (green carbon atoms). Hydrogen bonds are depicted as blue dashed lines. The inset at the top of the images shows the stereochemistry of the thiohemiketal formed by the nucleophilic attack of the catalytic Cys residue onto the α-keto group. A. Binding of **11a** to SARS-CoV M^pro^. The thiohemiketal is in the *R* configuration, with its oxygen accepting two hydrogen bonds from the oxyanion-hole amides of Gly143 and Cys145. The amide oxygen accepts an H-bond from His41. The side-chains of Ser144 and Arg188 have been omitted for clarity. B. The P2-benzyl substituent of **11a** cannot fully enter the S2 pocket of the HCoV-NL63 M^pro^, which is much smaller and has less plasticity than the corresponding pocket of SARS-CoV M^pro^ (cf. A). The benzyl therefore binds above the pocket in the view shown here; this is probably the reason for the total inactivity (IC_50_ > 50 μM) of compound **11a** against HCoV-NL63 M^pro^. The small size of the pocket is due to the replacement of the flexible Gln189 of the SARS-CoV M^pro^ by the more rigid Pro189 in this enzyme. The stereochemistry of the thiohemiketal is *R*. The side-chains of Ala143 and Gln188 have been omitted for clarity. C. Binding of **11a** to the CVB3 3C^pro^. The stereochemistry of the thiohemiketal is *S*, as the group accepts a hydrogen bond from His41, whereas the amide keto group accepts three H-bonds from the oxyanion hole (residues 145 - 147). The side-chain of Gln146 has been omitted for clarity. D. The crystal structure of **11f** in complex with HCoV-NL63 M^pro^ shows that this short (inactive) compound lacking a P3 residue has its P2-Boc group inserted into the S2 pocket of the protease. The stereochemistry of the thiohemiketal is *S*. The side-chains of Ala143 and Gln188 have been omitted for clarity. E. In contrast to P2 = benzyl in **11a**, the isobutyl group of **11n** is small and flexible enough to enter into the narrow S2 pocket of the HCoV-NL63 M^pro^. The thiohemiketal is in the *R* configuration. The side-chains of Ala143 and Gln188 have been omitted for clarity. F. In spite of its small size, the cyclopropylmethyl side-chain in the P2 position of **11s** can tightly bind to the S2 subsite of the SARS-CoV M^pro^, as this pocket exhibits pronounced plasticity due to the conformational flexibility of Gln189 (see also Fig. 4). The stereochemistry of the thiohemiketal is *S*. The side-chains of Ser144 and Arg188 have been omitted for clarity.

### P1’ and P3 substituents

The crystal structures indicated that the fits of the P1’ benzyl group of **11a** in the S1’ pocket and of the P3 cinnamoyl cap in the S3 subsite might be improved (see Fig. 3a-c). Compounds **11b** - **11e** and **11g** - **11l** were synthesized in an attempt to do so; however, none of them showed better inhibitory activity against the majority of the recombinant proteases, compared to the parent compound, **11a** (see Supplementary Results). To investigate whether the P3 residue of the inhibitor is dispensible, we synthesized compound **11f**, which only comprises P2 = Boc, P1 = GlnLactam, and P1’ = benzyl. **11f** was inactive against all purified proteases and in all replicons tested, but showed some activity against HRV2 in HeLa Rh cells (EC_50_ = 9.0 μM). A crystal structure of **11f** bound to HCoV-NL63 M^pro^ demonstrated that the P2-Boc group entered the S2 pocket (Fig. 3d). In conclusion, although there is probably room for further improvement, we decided to maintain the original design with P1’ = benzyl and P3 = cinnamoyl, and focussed on improving the P2 substituen.

### Properties of the S2 pockets of the target enzymes

The crystal structures of SARS-CoV M^pro^, HCoV-NL63 M^pro^, and CVB3 3C^pro^ in complex with **11a** revealed a fundamental difference between the S2 pockets of the coronavirus proteases and the enterovirus proteases: The cavities are covered by a “lid” in the former but are open to one side in the latter (Fig. 2,b-d). In SARS-CoV M^pro^, the lid is formed by the 3_10_ helix 46 - 51 and in HCoV-NL63 M^pro^ by the loop 43-48. Residues from the lid, in particular Met49 in the case of SARS-CoV M^pro^, can thus make hydrophobic interactions with the P2 substitutent of the inhibitor, whereas such interaction is missing in the enterovirus 3C^pro^s. In addition to the lid, the S2 pocket is lined by the “back-wall” (main-chain atoms of residues 186 and 188 and Cβ atom of Asp187), the side-walls (Gln189, His41), as well as the “floor” (Met165) in SARS-CoV M^pro^. In HCoV-NL63 M^pro^, the corresponding structural elements are main-chain atoms of residues 187 and 188 as well as the Cβ atom of Asp187 (back-wall), Pro189 and His41 (side-walls), and Ile165 (floor). Finally, in CVB3 3C^pro^, Arg39, Asn69, and Glu71 form the back-wall, residues 127-132 and His40 form the side-walls, and Val162 constitutes the floor.

In addition, the S2 pocket is of different size in the various proteases. The SARS-CoV enzyme features the largest S2 pocket, with a volume of 252 Å^3^ embraced by the residues (Gln189, His41) defining the side-walls of the pocket in the ligand-free enzyme, as calculated by using *Chimera*,^36^ followed by the CVB3 3C^pro^ S2 pocket with about 180 Å^3^ (space between Thr130 and His40). The HCoV-NL63 M^pro^ has by far the smallest S2 pocket of the three enzymes, with a free space of only 45 Å^3^ between Pro189 and His41, according to *Chimera*.

In agreement with these observations, a good fit is observed between the P2 benzyl group of **11a** and the S2 subsite of the SARS-CoV M^pro^ as well as that of the CVB3 3C^pro^ (Fig. 3a,c). In contrast, the crystal structure of the complex between **11a** and HCoV-NL63 M^pro^, against which the compound is inactive, demonstrates that the P2-benzyl group cannot fully enter the S2 pocket of the enzyme because of the restricted size of this site (Fig. 3b).

Thus, the properties of our target proteases with respect to the S2 pocket were defined at this point as “small” and “covered by a lid” for HCoV-NL63 M^pro^, “large” and “covered” for SARS-CoV M^pro^, and “large” and “open” for CVB3 3C^pro^. Through comparison with crystal structures of other proteases of the same virus genus (HCoV-229E M^pro^ for alphacoronaviruses^28^ (PDB entry 1P9S); HKU4 M^pro^ for betacoronaviruses (Ma, Xiao et al., unpublished; PDB entry 2YNA); and EV-A71 3C^pro^ for enteroviruses^30^ (PDB entry 3SGK), we ensured that our conclusions drawn from the template structures were valid for other family members as well.

To explore the sensitivity of the S2 pocket towards a polar substituent in the *para* position of the benzyl group, we synthesized compound **11m** carrying a 4-fluorobenzyl group in P2. This substitution abolished almost all activity against the SARS-CoV M^pro^ (IC_50_ > 50 μM), and the compound proved inactive against HCoV-NL63 M^pro^ as well, whereas IC_50_ values were 2.3 μM against the EV-A71 3C^pro^ and 8.7 μM against CVB3 3C^pro^. From this, we concluded that the introduction of the polar fluorine atom is not compatible with the geometry of the S2 pocket of SARS-CoV M^pro^, whereas the fluorine can accept a hydrogen bond from Arg39 in EV-A71 3C^pro^ (ref. 30) and probably also CVB3 3C^pro^. In SARS-CoV M^pro^, however, the carbonyl groups of residues 186 and 188 might lead to a repulsion of the fluorinated benzyl group.

### P2-alkyl substituents of varying sizes

As the P2-benzyl group of **11a** was apparently too large to fit into the S2 pocket of the HCoV-NL63 M^pro^, we replaced it by isobutyl in **11n**. This resulted in improved activities against SARS-CoV M^pro^ (IC_50_ = 0.33 μM) and in a very good activity against HCoV-NL63 M^pro^ (IC_50_ = 1.08 μM, compare with the inactive **11a**). For EV-A71 3C^pro^, however, the activity decreased to IC_50_ = 13.8 μM, different from CVB3 3C^pro^, where IC_50_ was 3.8 μM. Our interpretation of this result is that the smaller P2-isobutyl substituent of **11n** can still interact with the “lid” (in particular, Met49) of the SARS-CoV M^pro^ S2 site, but is unable to reach the “back-wall” of the EV-A71 3C^pro^ pocket and thus, in the absence of a “lid”, cannot generate sufficient enthalpy of binding. We will see from examples to follow that this trend persists among all inhibitors with a smaller P2 substituent: Even though the SARS-CoV M^pro^ S2 pocket has a larger volume than that of the enterovirus 3C^pro^, the enzyme can be efficiently inhibited by compounds carrying a small P2 residue that makes hydrophobic interactions with the lid (Met49) and floor (Met165) residues.

The EC_50_ of **11n** was >10 μM against the EV-A71 and CVB3 replicons, and even in the SARS-CoV replicon, the activity of **11n** was relatively weak (EC_50_ = 7.0 μM; Table 2). In agreement with the replicon data, **11n** proved inactive against EV-A71 in RD cells and showed limited activity against HRV2 or HRV14 in HeLa Rh cells (Table 3). Only the comparatively good activity (EC_50_ = 4.4 μM) against EV-D68 in HeLa Rh cells was unexpected. The activity of **11n** against HCoV 229E in Huh7 cells was good (EC_50_ = 0.6 μM), and against MERS-CoV in Huh7 cells, it was excellent, with EC_50_ = 0.0048 μM, while in Vero cells, the EC_50_ against MERS-CoV was as high as 9.2 μM. Similarly, the EC_50_ against SARS-CoV in Vero cells was 14.2 μM (Table 3).

We managed to obtain crystals of **11n** in complex with the M^pro^ of HCoV NL63 and found the P2 isobutyl group to be well embedded in the S2 pocket (Fig. 3e). This is not only a consequence of the smaller size of the isobutyl group compared to the benzyl group, but also of its larger conformational flexibility, which allows a better fit to the binding site.

When we replaced the P2-isobutyl residue of **11n** by n-butyl in **11o**, the activities were as follows: IC_50_ = 8.5 μM for SARS-CoV M^pro^, totally inactive (IC_50_ > 50 μM) against HCoV-NL63 M^pro^, IC_50_ = 3.2 μM for EV-A71 3C^pro^, and 5.2 μM for CVB3 3C^pro^. The decreased activity in case of SARS-CoV M^pro^ and the total inactivity against HCoV-NL63 M^pro^ indicate that the n-butyl chain is too long for the S2 pocket of these proteases, whereas the slight improvement against EV-A71 3C^pro^ and CVB3 3C^pro^ is probably a consequence of the extra space that is available to long and flexible substituents because of the lack of a lid covering the enterovirus 3C^pro^ pocket.

As the n-butyl substituent in P2 of **11o** was obviously too long, we next synthesized a derivative with the shorter propargyl (ethinylmethyl) as the P2 residue (compound **11p**). This led to very mediocre activities against all tested proteases. Using cyclopropyl as the P2 residue (compound **11q**), the IC_50_ values were even higher against most of the proteases tested. Obviously, the P2 side-chain requires a methylene group in the β-position in order to provide the necessary flexibility for the substituent to be embedded in the S2 pocket.

### Modifying ring size and flexibility of P2-cycloalkylmethyl substituents

Having realized that in addition to size, flexibility of the P2 substituent may be an important factor influencing inhibitory activity, we introduced flexibility into the phenyl ring of **11a** by reducing it. The cyclohexylmethyl derivative **11r** exhibited IC_50_ = 0.7 μM against SARS-CoV M^pro^, 12.3 μM against HCoV-NL63 M^pro^, 1.7 μM against EV-A71 3C^pro^, and 0.9 μM against CVB3 3C^pro^. Thus, the replacement of the phenyl group by the cyclohexyl group led to a significant improvement of the inhibitory activity against the recombinant SARS-CoV M^pro^ and to a dramatic improvement in case of CVB3 3C^pro^. Even for the HCoV-NL63 M^pro^, against which **11a** was completely inactive, greatly improved albeit still weak activity was observed (Table 1). In the viral replicons, **11r** performed very well, with EC_50_ = 0.8 - 0.9 μM for the EV-A71 replicon, 0.45 μM for CVB3, and 1.4 μM for SARS-CoV (Table 2). In the virus-infected cell culture assays (Table 3), **11r** exhibited EC_50_ = 3.7 μM against EV-A71 in RD cells and EC_50_ = 0.48 - 0.7 μM against EV-D68, HRV2, and HRV14 in HeLa cells. Against HCoV 229E in Huh7 cells, the EC_50_ was surprisingly low (1.8 μM). Interestingly, the compound proved extremely potent against MERS-CoV in Huh7 cells, with EC_50_ = 0.0004 μM (400 picomolar!). Even in Vero cells, EC_50_ against MERS-CoV was 5 μM, and the EC_50_ against SARS-CoV in Vero E6 cells was 1.8 - 2.1 μM, i.e. the best activity we have seen for an M^pro^ inhibitor against SARS-CoV in this type of cells. The therapeutic index (CC_50_/EC_50_) of **11r** against EV-D68, HRV2, and HRV14 was >15 in HeLa Rh cells as well as against CVB3 in Huh-T7 cells, but only ∼5 for EV-A71 in RD cells.

At this point, we decided to systematically vary the size of the ring system in P2. The next substituent to be tried was cyclopropylmethyl (compound **11s**, which showed good activities against SARS-CoV M^pro^ (IC_50_ = 0.24 μM) and HCoV-NL63 M^pro^ (1.4 μM), but poor values against EV-A71 3C^pro^ (IC_50_ = 18.5 μM) and CVB3 3C^pro^ (IC_50_ = 4.3 μM) (Table 1). **11s** was shown to inhibit the SARS-CoV replicon with an EC_50_ of about 2 μM, whereas activity against the EV-A71 and CVB3 replicons was poor (EC_50_ values > 20 μM) (Table 2). The replicon results were mirrored by the antiviral activity of **11s** in enter-ovirus-infected cells (Table 3), which was weak or very weak. By contrast, the compound inhibited HCoV 229E and MERS-CoV in Huh7 cells with EC_50_ of 1.3 and 0.08 μM, respectively. The activity against the latter virus in Vero cells was poor (EC_50_ ∼11 μM), and so was the anti-SARS-CoV activity in Vero E6 cells (Table 3).

We next analyzed the crystal structure of the complex between SARS-CoV M^pro^ and compound **11s** (Fig. 3f). The cyclopropylmethyl substituent was found to be incorporated deeply into the S2 pocket, making hydrophobic interactions with Met49 (the lid), Met165 (the floor), and the Cβ of Asp187 (the back-wall). In spite of the small size of the P2 substituent, this is possible because the S2 pocket of SARS-CoV M^pro^ is flexible enough to contract and enclose the P2 moiety tightly. This plasticity is expressed in a conformational change of residue Gln189, both in the main chain and in the side-chain. The main-chain conformational change is connected with a flip of the peptide between Gln189 and Thr190. The χ1 torsion angle of the Gln189 side-chain changes from roughly antiperiplanar (*ap*) to (-)-synclinal (*-sc*) (Fig. 4). The conformational variability of Gln189 has been noted before, both in Molecular Dynamics simulations^26^ and in other crystal structures.^37^ As a consequence of these changes, the side-chain oxygen of Gln189 can accept a 2.54-Å hydrogen bond from the main-chain NH of the P2 residue in the **11s** complex (see Fig. 4). The affinity of **11s** for the S2 pocket of HCoV-NL63 M^pro^ is good because of an almost ideal match of size and not requiring conformational changes, which this enzyme would not be able to undergo because of the replacement of the flexible Gln189 by the more rigid Pro. On the other hand, docking of the same compound into the crystal structure of the CVB3 3C^pro^ revealed that the cyclopropylmethyl moiety was probably unable to generate sufficient Free Energy of binding because of the missing lid and the large size of the S2 pocket in the enterovirus 3C^pro^, thereby explaining the poor inhibitory activity of **11s** against these targets.

**Figure 4:**
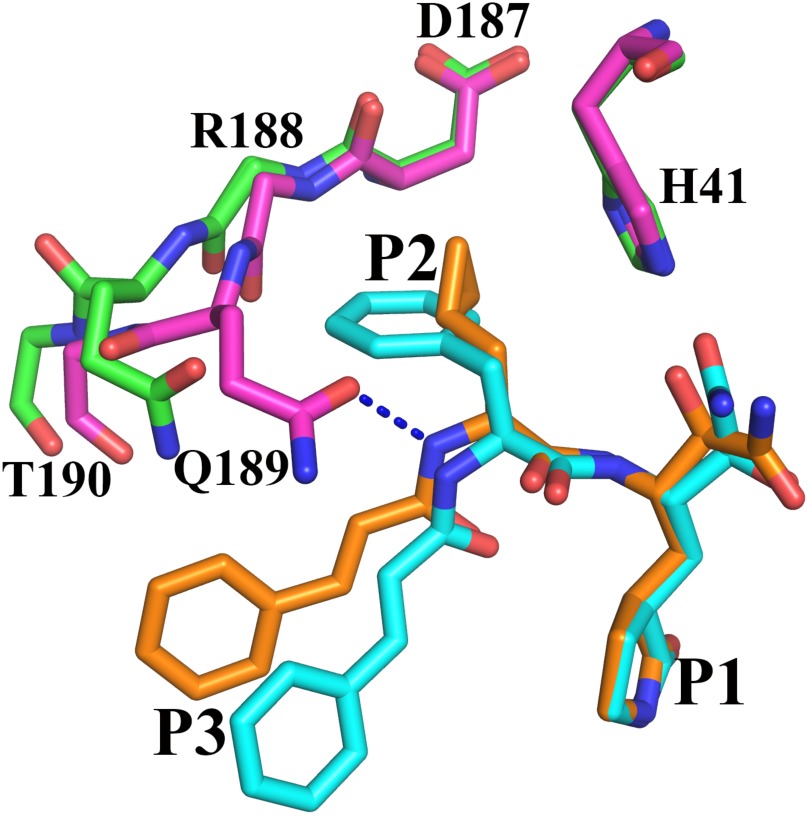
A pronounced plasticity of the S2 pocket of SARS-CoV Mpro is revealed by a comparison of the geometry of the subsite in the complexes with **11a** (P2 = benzyl; inhibitor cyan, protein green) and **11s** (P2 = cyclopropylmethyl; inhibitor orange, protein pink). The main differences here concern the main chain around Gln189 (note the flip of the 189 - 190 peptide bond) as well as the side-chain of this flexible residue, conformational change of which allows the S2 pocket to “shrink” and adapt to the small size of the P2-substituent in **11s**. This change also enables the formation of a hydrogen bond between the main-chain amide of the P2 residue and the side-chain oxygen of Gln189. The side-chains of Arg188 and Thr190 as well as the P1’ substituent of the inhibitors have been omitted for clarity.

We next introduced cyclobutylmethyl in the P2 position (compound **11t**) and obtained the following results: IC_50_ = 1.4 μM for SARS-CoV M^pro^, 3.4 μM for HCoV-NL63 M^pro^, 10.8 μM for EV-A71 3C^pro^, and 4.8 μM for CVB3 3C^pro^ (Table 1). Experiments with the viral replicons confirmed this trend, although the EC_50_ value for SARS-CoV (6.8 μM) was surprisingly high (Table 2). In Huh7 cells infected with MERS-CoV, this compound exhibited EC_50_ = 0.1 μM (but 9.8 μM in Vero cells), whereas EC_50_ was 7.0 μM against SARS-CoV in Vero E6 cells. The compound was largely inactive against EV-A71 in RD cells and inhibited the replication of the two HRV subtypes tested (in HeLa Rh cells) with EC_50_ values of ∼4 μM. The CC_50_ of **11t** in HeLa cells was 65 μM, i.e. the therapeutic index was well above 15 (Table 3).

Obviously, this substituent was still a bit too small for the enterovirus proteases, so as the next step, we tested P2 = cyclopentylmethyl (compound **11u**). This turned out to be the one compound with acceptable IC_50_ values against *all* tested enzymes: 1.3 μM against SARS-CoV M^pro^, 5.4 μM against HCoV-NL63 M^pro^, 4.7 μM against EV-A71 3C^pro^, and 1.9 μM against CVB3 3C^pro^ (Table 1). The activity against the replicons was between 3.6 and 4.9 μM (Table 2). In Huh7 cells infected with HCoV 229E or MERS-CoV, **11u** showed EC_50_ = 2.5 or 0.03 μM (11.1 μM for MERS-CoV in Vero cells), while EC_50_ was 4.9 μM against SARS-CoV in Vero E6 cells (Table 3).

**11u** appeared so far the best compromise compound, yet for each of the individual viral enzymes, the following compounds proved superior: P2 = cyclopropylmethyl (compound **11s**) for SARS-CoV M^pro^, P2 = isobutyl (compound **11n**) and P2 = cyclopropylmethyl (**11s**) for HCoV-NL63 M^pro^, P2 = benzyl (**11a)** or cyclohexylmethyl (**11r)** for EV-A71 3C^pro^, and **11r** for CVB3 3C^pro^. In other words, the nearly equipotent **11u** is indeed a compromise. Therefore, in view of the surprisingly good antiviral activity of **11r** against HCoV 229E in Huh7 cells, we relaxed the condition that the universal inhibitor should show good activity against the recombinant HCoV-NL63 M^pro^, and selected **11r** (P2 = cyclohexylmethyl) as the lead compound for further development. This compound exhibited submicromolar IC_50_ values against CVB3 3C^pro^ and SARS-CoV M^pro^, and IC_50_ = 1.7 μM against EV-A71 3C^pro^ (Table 1), as well as similarly low EC_50_ values in the replicons of these viruses (Table 2). In Huh7 cells infected with MERS-CoV, the performance of this compound was excellent, with EC_50_ = 0.0004 µM, and even against HCoV 229E in Huh7 cells and SARS-CoV in Vero E6 cells, EC_50_ values of 1.8 and 2.1 μM, respectively, were observed (Table 3). Also in enterovirus-infected cell culture, the compound performed well, with EC_50_ values of 0.7 μM or below against HRV2, HRV14, and EV-D68 in HeLa (Rh) cells and selectivity values >15. The only concern is the activity of the compound against EV-A71 in RD cells, for which the EC_50_ value was 3.7 μM, resulting in too low a therapeutic index. On the other hand, only weak toxicity was detected for **11r** in Vero or Huh-T7 cells. Preliminary pharmacokinetics tests with the compound in mice did not indicate a toxicity problem (to be published elsewhere).

## DISCUSSION

We describe here the structure-based design, the synthesis, and the assessment of capped dipeptide α-ketoamides that target the main protease of alpha- or betacoronaviruses as well as the 3C protease of enteroviruses. Through crystallographic analyses of a total of six inhibitor complexes of three different proteases in this study, we found the α-ketoamide warhead (-CO-CO-NH-) to be sterically more versatile than other warheads such as Michael acceptors (-CH=CH-CO-) and aldehydes (-CH=O), because it features two acceptors for hydrogen bonds from the protein, namely the α-keto oxygen and the amide oxygen, whereas the other warheads have only one such acceptor. In the various complexes, the hydroxy group (or oxyanion) of the thiohemiketal that is formed by the nucleophilic attack of the active-site cysteine residue onto the α-keto carbon, can accept one or two hydrogen bonds from the main-chain amides of the oxyanion hole. In addition, the amide oxygen of the inhibitor accepts a hydrogen bond from the catalytic His side-chain. Alternatively, the thiohemiketal can interact with the catalytic His residue and the amide oxygen with the main-chain amides of the oxyanion hole. Depending on the exact interaction, the stereochemistry at the thiohemiketal C atom would be different. We have previously observed a similar difference in case of aldehyde inhibitors, where the single interaction point — the oxyanion of the thiohemiacetal — can accept a hydrogen bond either from the oxyanion hole or from the catalytic His side-chain,^37^ resulting in different stereochemistry of the thiohemiacetal carbon. Both α-ketoamides and aldehydes react reversibly with the catalytic nucleophile of proteases, whereas Michael acceptors form irreversible adducts.

In addition to better matching the H-bonding donor/acceptor properties of the catalytic center through offering two hydrogen-bond acceptors instead of one, α-ketoamides have another big advantage over aldehydes and α,β-unsaturated esters (Michael acceptors) in that they allow easy extension of the inhibitors to probe the primed specificity subsites beyond S1’, although this has so far rarely been explored (e.g., ref. 38 in case of calpain).

The most prominent α-ketoamide drugs are probably telaprivir and boceprivir, peptidomimetic inhibitors of the HCV NS3/4A protease,^39,40^ which have helped revolutionize the treatment of chronic HCV infections. For viral cysteine proteases, α-ketoamides have only occasionally been described as inhibitors and few systematic studies have been carried out.

A number of capped dipeptidyl α-ketoamides have been described as inhibitors of the norovirus 3C-like protease.^41^ These were optimized with respect to their P1’ substituent, whereas P2 was isobutyl in most cases and occasionally benzyl. The former displayed IC_50_ values one order of magnitude lower than the latter, indicating that the S2 pocket of the norovirus 3CL protease is fairly small. Although we did not include the norovirus 3CL^pro^ in our study, expanding the target range of our inhibitors to norovirus is probably a realistic undertaking.

While our study was underway, Zeng et al.^42^ published a series of α-ketoamides as inhibitors of the EV-A71 3C^pro^. These authors mainly studied the structure-activity relationships of the P1’ residue and found small alkyl substituents to be superior to larger ones. Interestingly, they also reported that a six-membered δ-lactam in the P1 position led to 2 - 3 times higher activities, compared to the five-membered γ-lactam. At the same time, Kim et al.^43^ described a series of five α-ketoamides with P1’ = cyclopropyl that showed submicromolar activity against EV-D68 and two HRV strains.

Occasionally, individual α-ketoamides have been reported in the literature as inhibitors of both the enterovirus 3C protease and the coronavirus main protease. A single capped dipeptidyl α-ketoamide, Cbz-Leu-GlnLactam-CO-CO-NH-iPr, was described that inhibited the recombinant transmissible gastroenteritis virus (TGEV) and SARS-CoV M^pro^s as well as human rhinovirus and poliovirus 3C^pro^s in the one-digit micromolar range.^44^ Coded GC-375, this compound showed poor activity in cell culture against EV-A71 though (EC_50_ = 15.2 μM), probably because P2 was isobutyl. As we have shown here, an isobutyl side-chain in the P2 position of the inhibitors is too small to completely fill the S2 pocket of the EV-A71 3C^pro^ and the CVB3 3Cpro.

Among a series of aldehydes, Prior et al.^45^ described the capped tripeptidyl α-ketoamide Cbz-1-naphthylalanine-Leu-GlnLactam-CO-CO-NH-iPr, which showed IC_50_ values in the 3-digit nanomolar range against HRV 3C^pro^ and SARS-CoV M^pro^, as well as EC_50_ values of 0.03 μM against HRV18 and 0.5 μM against HCoV 229E in cell culture. No optimization of this compound was performed and no toxicity data have been reported.

For compounds with warheads other than α-ketoamides, *in-vitro* activity against both corona- and enteroviruses has also occasionally been reported. Lee et al.^46^ described three peptidyl Michael acceptors that displayed inhibitory activity against the M^pro^s of SARS-CoV and HCoV 229E as well as against the 3C^pro^ of CVB3. These inhibitors had an IC_50_ 10 - 20 times higher for the CVB3 enzyme, compared to SARS-CoV M^pro^. P2 was invariably isobutyl (leucine) in these compounds, suggesting that further improvement might be possible.

In addition to Michael acceptors, peptide aldehydes have also been used to explore the inhibition of coronavirus M^pro^s as well as enterovirus 3C^pro^s. Kim et al.^44^ reported a dipeptidyl aldehyde and its bisulfite adduct, both of which exhibited good inhibitory activities against the isolated 3C proteases of human rhinovirus and poliovirus as well as against the 3C-like proteases of a number of coronaviruses, but antiviral activities in cell culture against EV-A71 were poor (EC_50_ >10 μM), again most probably due to P2 being isobutyl (leucine).

In our series of compounds, we used P1 = GlnLactam (γ-lactam) throughout, because this substituent has proven to be an excellent surrogate for glutamine.^29,32^ While we made some efforts to optimize the P1’ residue of the compounds as well as the N-cap (P3), we mainly focussed on optimization of the P2 substituent. In nearly all studies aiming at discovering peptidomimetic inhibitors of coronavirus M^pro^s, P2 is invariably isobutyl (leucine), and this residue has also been used in the efforts to design compounds that would inhibit enterovirus 3C^pro^s as well (see above). From crystal structures of our early lead compound, **11a** (cinnamoyl-Phe-GlnLactam-CO-CO-NH-Bz), in complex with the M^pro^s of HCoV NL63 (as representative of the alphacoronavirus proteases) and SARS-CoV (beta-CoV) as well as the 3C^pro^ of Coxsackievirus B3 (enterovirus proteases), we found that the S2 pocket has fundamentally different shapes in these enzymes. In the SARS-CoV M^pro^, the S2 subsite is a deep hydrophobic pocket that is truly three-dimensional in shape: the “walls” of the groove are formed by the polypeptide main chain around residues 186 - 188 as well as by the side-chains of His41 (of the catalytic dyad) and Gln189, whereas the “floor” is formed by Met165 and the “lid” by residues 45 - 51, in particular Met49. The two methionines provide important interaction points for the P2 substituents of inhibitors; while these interactions are mostly hydrophobic in character, we have previously described the surprising observation of the carboxylate of an aspartic residue in P2 that made polar interactions with the sulfur atoms of these methionines.^37^ Because the pocket offers so many opportunities for interaction and features a pronounced plasticity, P2 substituents such as isobutyl (from Leu), which are too small to fill the pocket entirely, can still generate sufficient binding enthalpy. Accordingly, the S2 pocket of SARS-CoV M^pro^ is the most tolerant among the three enzymes investigated here, in terms of versatility of the P2-substituents accepted.

In the S2 pocket of the HCoV-NL63 M^pro^, Gln189 is replaced by proline and this change is accompanied by a significant loss of flexibility; whereas the side-chain of Gln189 of SARS-CoV M^pro^ is found to accommodate its conformation according to the steric requirements of the P2 substituent, the proline is less flexible, leading to a much smaller space at the entrance to the pocket. As a consequence, a P2-benzyl substituent is hindered from penetrating deeply into the pocket, whereas the smaller and more flexible isobutyl group of P2-Leu is not.

Finally, in the 3C^pro^s of EV-A71 and CVB3, the S2 pocket lacks a lid, i.e. it is open to one side. As a consequence, it offers less interaction points for P2 substituents of inhibitors, so that such substituents must reach the “back-wall” of the pocket (formed by Arg39, Asn69, Glu71) in order to create sufficient binding energy. Hence, large aromatic substituents such as benzyl are favored by the enterovirus 3C^pro^s.

When we introduced a fluoro substituent in the *para* position of the P2-benzyl group of our lead compound, **11a**, we observed good activity against the enterovirus 3C^pro^s but complete inactivity against the coronavirus M^pro^s (see Table 1, compound **11m**). This is easily explained on the basis of the crystal structures: In the enterovirus 3C^pro^s, the fluorine can accept a hydrogen bond from Arg39 (ref. 30), whereas in the coronavirus M^pro^s, there would be electrostatic repulsion from the main-chain carbonyls of residues 186 and 188. In agreement with this, rupintrivir (which has P2 = *p*-fluorobenzyl) is a good inhibitor of the enteroviral 3C^pro^s,^46^ but not of the coronaviral main proteases, as we predicted earlier.^28^

In this structure-based inhibitor optimization study, we achieved major improvements over our original lead compound, **11a**, by systematically varying the size and the flexibility of the P2-substituent. The compound presenting so far the best compromise between the different requirements of the S2 pockets (SARS-CoV M^pro^: large and covered, HCoV-NL63 M^pro^: small and covered, and CVB3 3C^pro^: large and open) is **11u** (P2 = cyclopentylmethyl), which has satisfactory broad-spectrum activity against all proteases tested. However, with regard to its antiviral activities in cell cultures, it is inferior to **11r** (P2 = cyclohexylmethyl). The latter compound exhibits very good inhibitory activity against the SARS-CoV M^pro^ as well as the enterovirus 3C^pro^, and its performance in the SARS-CoV and enterovirus replicons is convincing. Being in the low micromolar range (EV-A71, CVB3), the data for the antiviral activity in cell culture for **11r** correlate well with the inhibitory power of the compound against the recombinant proteases as well as in the replicon-based assays. This is not true, though, for the surprisingly good *in-cellulo* activity of **11r** against HCoV 229E in Huh7 cells. Also, the correlation does not seem to hold for LLC-MK2 and CaCo2 cells. We tested the antiviral activity of many of our compounds against HCoV NL63 in these two cell types and found that all of them had low- or submicromolar EC_50_ values against this virus in LLC-MK2 cells but were largely inactive in CaCo2 cells (not shown). Furthermore, **11r** and all other compounds that we synthesized are inactive (EC_50_ > 87 μM) against CVB3 in Vero cells (not shown), but exhibit good to excellent activities against the same virus in Huh-T7 cells. We have previously observed similar poor antiviral activities in Vero cells not only for α-ketoamides, but also for Michael acceptors (Zhu et al., unpublished work). A similar cell-type dependence is seen for the antiviral activity of **11r** against MERS-CoV and SARS-CoV. Whereas the inhibitor exhibits excellent activity against MERS-CoV when Huh7 cells are the host cells (400 pM), the inhibitory activity is weaker by a factor of up to 12,500 when Vero cells are used (EC_50_ = 5 μM). On the other hand, **11r** exhibits excellent anti-MERS-CoV activity in human Calu3 lung cells, i.e. in the primary target cells where the compound will have to act in a therapeutic setting (A. Kupke, personal communication). As we tested antiviral activity against SARS-CoV exclusively in Vero cells, the EC_50_ values determined for our compounds against this virus are in the one-digit micromolar range or higher; the best is again compound **11r** with EC_50_ = 2.1 μM. Interestingly, the relatively weaker activity (or even inactivity) of our inhibitors against RNA viruses in Vero cells was observed independently in the virology laboratories in Leuven and in Leiden. It is thus unlikely that the lack of activity in Vero cells is related to problems with the experimental set-up. In preliminary experiments, we replaced the P3 cinnamoyl group of **11r** by the fluorophor coumaryl and found by fluorescence microscopy that much more inhibitor appeared to accumulate in Huh7 cells compared to Vero cells (D.L., R.H. & Irina Majoul, unpublished).

Regardless of which cell system is the most suitable one for the testing of peptidomimetic antiviral compounds, we next plan to test **11r** in small-animal models for MERS and for Coxsackievirus-induced pancreatitis. In parallel, we aim to refine the experiments to quantify the accumulation of peptidomimetic protease inhibitors in different host-cell types, in the hope to find an explanation for the observed cell-type dependencies.

## CONCLUSIONS

This work demonstrates the power of structure-based approaches in the design of broad-spectrum antiviral compounds with roughly equipotent activity against coronaviruses and enteroviruses. We observed a good correlation between the inhibitory activity of the designed compounds against the isolated proteases, in viral replicons, and in virus-infected Huh7 cells. One of the compounds (**11r**) exhibits excellent anti-MERS-CoV activity in virus-infected Huh7 cells. Because of the high similarity between the main proteases of SARS-CoV and the novel BetaCoV/Wuhan/2019, we expect **11r** to exhibit good antiviral activity against the new coronavirus as well.

## EXPERIMENTAL SECTION

### Crystallization and X-ray structure determination of complexes between viral proteases and α-ketoamides

#### Crystallization

The recombinant production and purification of SARS-CoV M^pro^ with authentic N and C termini was described in detail previously.^48,49^ Using an Amicon YM10 membrane (EMD Millipore), the purified SARS-CoV M^pro^ was concentrated to 21 mg·mL^-1^ in buffer A (20 mM Tris-HCl, 150 mM NaCl, 1 mM DTT, 1 mM EDTA, pH 7.5). Crystallization was performed by equilibrating 1 μL protein (mixed with 1 μL precipitant solution) against 500 μl reservoir containing 6 – 8% polyethylene glycol (PEG) 6,000, 0.1 M MES (pH 6.0), at 20°C using the vapor diffusion sitting-drop method. Compounds **11a** and **11s** were dissolved in 100% DMSO at 50 mM and 200 mM stock concentrations, respectively. A crystal of the free enzyme was soaked in cryo-protectant buffer containing 20% MPD, 6% PEG 6,000, 0.1 M MES, 7.5 mM **11a**, pH 6.0, for 2 h at 20°C. Another set of free enzyme crystals was soaked in another cryo-protectant buffer with 6% PEG 6,000, 5% MPD, 0.1 M MES, 15% glycerol, 10 mM **11s**, pH 6.0, for 2 h. Subsequently, crystals were fished and flash-cooled in liquid nitrogen prior to data collection.

Crystals of HCoV-NL63 M^pro^ with **11a** were obtained using cocrystallization. The concentrated HCoV-NL63 M^pro^ (45 mg·mL^-1^) was incubated with 5 mM **11a** for 4 h at 20°C, followed by setting up crystallization using the vapor diffusion sitting-drop method at 20°C with equilibration of 1 μL protein (mixed with 1 μL mother liquor) against 500 μL reservoir composed of 0.1 M lithium sulfate monohydrate, 0.1 M sodium citrate tribasic dihydrate, 25% PEG 1,000, PH 6.0. The crystals were protected by a cryo-buffer containing 0.1 M lithium sulfate monohydrate, 0.1 M sodium citrate tribasic dihydrate, 25% PEG 1,000, 15% glycerol, 2 mM **11a**, pH 6.0 and flash-cooled in liquid nitrogen.

Crystals of HCoV-NL63 M^pro^ with **11n** or **11f** were generated by using the soaking method. Several free-enzyme crystals were soaked in cryo-protectant buffer containing 0.1 M lithium sulfate monohydrate, 0.1 M sodium citrate tribasic dihydrate, 25% PEG 1,000, 15% glycerol, 5 mM **11n (**or **11f**), pH 6.0. Subsequently, the soaked crystals were flash-cooled in liquid nitrogen.

Freshly prepared CVB3 3C^pro^ at a concentration of 21.8 mg·mL^-1^ was incubated with 5 mM **11a** pre-dissolved in 100% DMSO at room tempature for 1 h. Some white precipitate appeared in the mixture. Afterwards, the sample was centrifuged at 13,000 × *g* for 20 min at 4 °C. The supernatant was subjected to crystallization trials using the following, commercially available kits: Sigma™ (Sigma-Aldrich), Index™, and PEG Rx™ (Hampton Research). Single rod-like crystals were detected both from the Index™ screen, under the condition of 0.1 M MgCl_2_ hexahydrate, 0.1 M Bis-Tris, 25% PEG 3,350, pH 5.5, and from the Sigma™ screen at 0.2 M Li_2_SO_4_, 0.1 M Tris-HCl, 30% PEG 4,000, pH 8.5. Crystal optimization was performed by using the vapor-diffusion sitting-drop method, with 1 μL CVB3 3C^pro^-inhibitor complex mixed with 1 μL precipitant solution, and equilibration against 500 μL reservoir containing 0.1 M Tris-HCl, 0.2 M MgCl_2_, pH 8.5, and PEG 3,350 varied from 22% to 27%. Another optimization screen was also performed against a different reservoir, 0.1 M Tris-HCl, 0.2 M MgCl_2_, pH range from 7.5 to 8.5, and PEG 4,000 varied from 24% to 34%. Crystals were fished from different drops and protected by cryo-protectant solution consisting of the mother liquor and 10% glycerol. Subsequently, the crystals were flash-cooled with liquid nitrogen.

#### Diffraction data collection, structure elucidation and refinement

Diffraction data from the crystal of the SARS-CoV M^pro^ in complex with **11a** were collected at 100 K at synchrotron beamline PXI-X06SA (PSI, Villigen, Switzerland) using a Pilatus 6M detector (DECTRIS). A diffraction data set from the SARS-CoV M^pro^ crystal with compound **11s** was collected at 100 K at beamline P11 of PETRA III (DESY, Hamburg, Germany), using the same type of detector. All diffraction data sets of HCoV-NL63 M^pro^ complex structures and of the complex of CVB3 3C^pro^ with **11a** were collected at synchrotron beamline BL14.2 of BESSY (Berlin, Germany), using an MX225 CCD detector (Rayonics). All data sets were processed by the program XDSAPP and scaled by SCALA from the CCP4 suite.^50-52^ The structure of SARS-CoV M^pro^ with **11a** was determined by molecular replacement with the structure of the complex between SARS-CoV M^pro^ and **SG85** (PDB entry 3TNT; Zhu et al., unpublished) as search model, employing the MOLREP program (also from the CCP4 suite).^52,53^ The complex structures of HCoV-NL63 M^pro^ with **11a, 11f**, and **11n** were also determined with MOLREP, using as a search model the structure of the free enzyme determined by us (LZ et al., unpublished). The complex structure between CVB3 3C^pro^ and **11a** was determined based on the search model of the free-enzyme structure (PDB entry 3ZYD; Tan et al., unpublished). Geometric restraints for the compounds **11a, 11f, 11n**, and **11s** were generated by using JLIGAND^52,54^ and built into F_o_-F_c_ difference density using the COOT software.^55^ Refinement of the structures was performed with REFMAC version 5.8.0131 (ref. 52,56,57).

#### Inhibitory activity assay of alpha-ketoamides

A buffer containing 20 mM Tris-HCl, 100 mM NaCl, 1 mM EDTA, 1 mM DTT, pH 7.3, was used for all the enzymatic assays. Two substrates with the cleavage sites of M^pro^ and 3C^pro^, respectively (indicated by the arrow, ↓), Dabcyl-KTSAVLQ↓SGFRKM-E(Edans)-NH_2_ and Dabcyl-KEALFQ↓GPPQF-E(Edans)-NH_2_ (95% purity; Biosyntan), were employed in the fluorescence resonance energy transfer (FRET)-based cleavage assay, using a 96-well microtiter plate. The dequenching of the Edans fluorescence due to the cleavage of the substrate as catalyzed by the proteases was monitored at 460 nm with excitation at 360 nm, using a Flx800 fluorescence spectrophotometer (BioTek). Curves of relative fluorescence units (RFU) against substrate concentration were linear for all substrates up to beyond 50 μM, indicating a minimal influence of the inner-filter effect. Stock solutions of the compounds were prepared by dissolving them in 100% DMSO. The UV absorption of **11a** was found to be negligible at λ = 360 nm, so that no interference with the FRET signal through the inner-filter effect was to be expected. For the determination of the IC_50_, different proteases at a specified final concentration (0.5 μM SARS-CoV or HCoV-NL63 M^pro^, 2 μM CVB3 3C^pro^, 3 μM EV-A71 3C^pro^) were separately incubated with the inhibitor at various concentrations (0 to 100 μM) in reaction buffer at 37°C for 10 min. Afterwards, the reaction was initiated by adding FRET peptide substrate at 20 μM final concentration (final volume: 50 μl). The IC_50_ value was determined by using the GraphPad Prism 6.0 software (GraphPad). Measurements of enzymatic activity were performed in triplicate and are presented as the mean ± standard deviations (SD).

Assessment of inhibitory activity of α-ketoamides using viral replicons and virus-infected cells

#### Cells and viruses

Hepatocellular carcinoma cells (Huh7; ref. 58) and their derivative constitutively expressing T7 RNA polymerase (Huh-T7; ref. 59) were grown in Dulbecco’s modified minimal essential medium (DMEM) supplemented with 2 mM glutamine, 100 U·mL^-1^ penicillin, 100 µg·mL^-1^ streptomycin sulfate, and fetal calf serum (10% in growth medium and 2% in maintenance medium). Huh-T7 cells were additionally supplemented with geneticin (G-418 sulfate, 400 µg·mL^-1^). Huh-T7 cells were used for the enteroviral replicons as well as for infection experiments with CVB3 strain Nancy.

For enterovirus (except CVB3) infection experiments, human rhabdomyosarcoma cells (RD; for EV-A71; BRCR strain) and HeLa Rh cells (for EV-D68 and human rhinoviruses) were grown in MEM Rega 3 medium supplemented with 1% sodium bicarbonate, 1% L-glutamine, and fetal calf serum (10% in growth medium and 2% in maintenance medium). For HCoV-229E (a kind gift from Volker Thiel (Bern, Switzerland)), culture and infection experiments were carried out as described.^60^ For MERS-CoV or SARS-CoV infection experiments, Vero, Vero E6, and Huh7 cells were cultured as described previously.^61,62^ Infection of Vero and Huh7 cells with MERS-CoV (strain EMC/2012) and SARS-CoV infection of Vero E6 cells (strain Frankfurt-1) at low multiplicity of infection (MOI) were done as described before.^61,63^ All work with live MERS-CoV and SARS-CoV was performed inside biosafety cabinets in biosafety level-3 facilities at Leiden University Medical Center, The Netherlands.

#### Viral replicons

The DNA-launched SARS-CoV replicon harbouring Renilla luciferase as reporter directly downstream of the SARS-CoV replicase polyprotein-coding sequence (pp1a, pp1ab, Urbani strain, acc. AY278741), in the context of a bacterial artificial chromosome (BAC) under the control of the CMV promoter, has been described previously (pBAC-REP-RLuc).^33^ Apart from the replicase polyprotein, the replicon encodes the following features: the 5’- and 3’-non-translated regions (NTR), a ribozyme (Rz), the bovine growth hormone sequence, and structural protein N.

Subgenomic replicons of CVB3 (pT7-CVB3-FLuc^34^) and EV-A71 (pT7-EV71-RLuc) harbouring T7-controlled complete viral genomes, in which the P1 capsid-coding sequence was replaced by the Firefly (*Photinus pyralis*) or Renilla (*Renilla renifor)* luciferase gene, were generously provided by F. van Kuppeveld and B. Zhang, respectively. To prepare CVB3 and EV-A71 replicon RNA transcripts, plasmid DNAs were linearized by digestion with SalI or HindIII (New England Biolabs), respectively. Copy RNA transcripts were synthesized *in vitro* using linearized DNA templates, T7 RNA polymerase, and the T7 RiboMax™ Large-Scale RNA Production System (Promega) according to the manufacturer’s recommendations.

#### Transfection

Huh-T7 cells grown in 12-well plates to a confluency of 80% - 90% (2 - 3 × 10^5^ cells/well) were washed with 1 mL OptiMEM (Invitrogen) and transfected with 0.25 µg of the replication-competent replicon and Lipofectamin2000 or X-tremeGENE9 in 300 µl OptiMEM (final volume) as recommended by the manufacturer (Invitrogen or Roche, respectively). The transfection mixtures were incubated at 37°C for 4 to 5 h (Lipofectamin2000) or overnight (X-tremeGENE9), prior to being replaced with growth medium containing the compound under investigation. For RNA-launched transfection of enteroviral replicons, DMRIE-C was used as transfection reagent according to the manufacturer’s recommendations (Invitrogen). All experiments were done in triplicate or quadruplicate and the results are presented as mean values ± SD.

#### Testing for inhibitory activity of candidate compounds

Initially, we performed a quick assessment of the inhibitory activity of the candidate compounds towards the enteroviral and coronaviral replicons at a concentration of 40 μM in Huh-T7 cells. Compounds that were relatively powerful and non-toxic at this concentration, were assayed in a dose-dependent manner to estimate their half-maximal effective concentration (EC_50_) as well as their cytotoxicity (CC_50_), as described.^29^ In brief, different concentrations of α-ketoamides (40 μM in screening experiments or increasing concentration (0, 1.25, 2.5, 5, 10, 20, 40 μM) when determining the EC_50_) were added to growth medium of replicon-transfected Huh-T7 cells. Twenty-four hours later, the cells were washed with 1 mL phosphate-buffered saline (PBS or OPTIMEM, Invitrogen) and lysed in 0.15 mL Passive lysis buffer (Promega) at room temperature (RT) for 10 min. After freezing (−80°C) and thawing (RT), the cell debris was removed by centrifugation (16,000 × *g*, 1 min) and the supernatant (10 or 20 μl) was assayed for Firefly or Renilla luciferase activity (Promega or Biotrend Chemikalien) using an Anthos Lucy-3 luminescence plate reader (Anthos Microsystem).

#### Antiviral assay with infectious enteroviruses

The antiviral activity of the compounds was evaluated in a cytopathic effect (CPE) readout assay using MTS [3-(4,5-dimethylthiazol-2-yl)-5-(3-carboxymethoxyphenol)-2-(4-sulfophenyl)-2H-tetrazolium, inner-salt]-based assay. Briefly, 24 h prior to infection, cells were seeded in 96-well plates at a density of 2.5 × 10^4^ (RD cells) or of 1.7 × 10^4^ (HeLa Rh) per well in medium supplemented with 2% FCS. For HRV2 and HRV14 infection, the medium contained 30 mM MgCl_2_. The next day, serial dilutions of the compounds and virus inoculum were added. The read-out was performed 3 days post infection as follows: The medium was removed and 100 μl of 5% MTS in Phenol Red-free MEM was added to each well. Plates were incubated for 1 h at 37°C, then the optical density at 498 nm (OD_498_) of each well was measured by microtiter plate reader (Saffire^2^, Tecan). The OD values were converted to percentage of controls and the EC_50_ was calculated by logarithmic interpolation as the concentration of compound that results in a 50% protective effect against virus-induced CPE. For each condition, cell morphology was also evaluated microscopically.

#### Antiviral assays with SARS and MERS coronaviruses

Assays with MERS-CoV and SARS-CoV were performed as previously described.^61,63^ In brief, Huh7, Vero, or Vero E6 cells were seeded in 96-well plates at a density of 1 × 10^4^ (Huh7 and Vero E6) or 2 × 10^4^ cells (Vero) per well. After overnight growth, cells were treated with the indicated compound concentrations or DMSO (solvent control) and infected with an MOI of 0.005 (final volume 150 μl/well in Eagle’s minimal essential medium (EMEM) containing 2% FCS, 2 mM *L*-glutamine, and antibiotics). Huh7 cells were incubated for two days and Vero/VeroE6 cells for three days, and differences in cell viability caused by virus-induced CPE or by compound-specific side effects were analyzed using the CellTiter 96 AQ_ueous_ Non-Radioactive Cell Proliferation Assay (Promega), according to the manufacturer’s instructions. Absorbance at 490 nm (A_490_) was measured using a Berthold Mithras LB 940 96-well plate reader (Berthold). Cytotoxic effects caused by compound treatment alone were monitored in parallel plates containing mock-infected cells.

#### Antiviral assay with Human coronavirus 229E

For HCoV-229E infection experiments, 5×10^4^ Huh7 cells were infected in triplicate in 24-well plates in 100 µl DMEM at 0.1 pfu/mL. After 1.5 h incubation at 37°C, virus inocula were removed. Cells were washed with DMEM and complete DMEM (10% FCS, 1% Pen./Strep.) containing the desired concentration of inhibitors (0, 1, 2.5, 5, 10, 20, and 40 μM) were added. After 48 h, the supernatant was collected. Viral RNA was isolated using the Bioline ISOLATE II RNA Mini Kit (#BIO-52072) according to the manufacturer’s instructions and eluted in 30 μl RNase-free water. qPCR was performed using the Bioline SensiFAST Probe Hi-ROX One-Step Kit (# BIO-77001) in a Roche Light Cycler96. cDNA was synthesized at 48°C for 1800 sec and 95°C for 600 sec, followed by 45 cycles at 95°C for 15 sec and 60°C for 60 sec at a temperature ramp of 4.4°C/sec. qPCR primer sequences (adapted from ref. 64) were 229E-For: 5’-CTACAGATAGAAAAGTTGCTTT-3’, HCoV-229E-Rev: 5′-ggTCGTTTAGTTGAGAAAAGT -3′, and 229E-ZNA probe: 5’- 6- Fam-AGA (pdC)TT(pdU)G(pdU)GT(pdC)TA(pdC)T-ZNA-3-BHQ-1 -3’ (Metabion). Standard curves were prepared using serial dilutions of RNA isolated from virus stock. Data were analysed using GraphPad Prism 5.0; EC_50_ values were calculated based on a 4-parameter logistic statistics equation. In parallel to the qPCR assays with inhibitors, cell viability assays were performed using Alamar-Blue™ Cell Viability Reagent (ThermoFisher) according to the manufacturer’s instruction. CC_50_ values were calculated using inhibitor versus normalized response statistics equation by including proper controls (no inhibitor and 1% Triton-X-100-treated cells).

#### Determination of the cell toxicity of candidate compounds

The CellTiter 96Aqueous One Solution Cell Proliferation Assay (MTS test, Promega), the CellTiter Glo assay kit (Promega), the Non-Destructive Cytotoxicity Bio-Assay (ToxiLight (measuring the release of adenylate kinase from damaged cells), Lonza Rockland), or the AlamarBlue™ Cell Viability Reagent (ThermoFisher) were used to determine the cytotoxic effect of compounds towards host cells according to the manufacturers’ recommendations.^29,65^

### Chemical synthesis of α-ketoamides

#### General procedure

Reagents were purchased from commercial sources and used without purification. HSGF 254 (0.15 - 0.2 mm thickness) was used for analytical thin-layer chromatography (TLC). All products were characterized by their NMR and MS spectra. ^1^H NMR spectra were recorded on 300-MHz, 400-MHz, or 500-MHz instruments. Chemical shifts were reported in parts per million (ppm, δ) down-field from tetramethylsilane. Proton coupling patterns were described as singlet (s), doublet (d), triplet (t), quartet (q), multiplet (m), and broad (br). Mass spectra were recorded using a Bruker ESI ion-trap HCT Ultra. HPLC spectra were recorded by LC20A or LC10A (Shimadzu Corporation) with Shim-pack GIST C18 (5 µm, 4.6×150mm) with three solvent systems (methanol/water, methanol/ 0.1% HCOOH in water or methanol/0.1% ammonia in water). Purity was determined by reversed-phase HPLC and was ≥95% for all compounds tested biologically.

Synthesis of (2*S*,4*R*)-dimethyl 2-(*tert*-butoxycarbonylamino)-4-(cyanomethyl) pentanedioate (1).

To a solution of *N*-Boc-*L*-glutamic acid dimethyl ester (6.0 g, 21.8 mmol) in THF (60 mL) was added dropwise a solution of lithium bis(trimethylsilyl)amide (LHMDS) in THF (47 mL, 1 M) at -78°C under nitrogen. The resulting dark mixture was stirred at -78°C. Meanwhile, bromoacetonitrile (1.62 mL, 23.3 mmol) was added dropwise to the dianion solution over a period of 1 h while keeping the temperature below -70°C. The reaction mixture was stirred at -78°C for additional 2 h. After the consumption of the reactant was confirmed by TLC analysis, the reaction was quenched by methanol (3 mL), and acetic acid (3 mL) in precooled THF (20 mL) was added. After stirring for 30 min, the cooling bath was removed. The reaction mixture was allowed to warm up to room temperature and then poured into brine (40 mL). The organic layer was concentrated and purified by flash column chromatography (petroleum ether/ethyl acetate = 4/1) to give product **1** (4.92 g, 72%) as colorless oil. ^1^H NMR (CDCl_3_, 400 MHz) δ 5.23 (1H, d, *J* = 9.0 Hz), 4.43-4.36 (1H, m), 3.77(1H, s), 3.76 (1H, s), 2.89-2.69 (3H, m), 2.20-2.14 (2H, m), 1.45 (9H, s). ESI-MS (*m*/*z*): 315 (M + H)^+^.

Synthesis of (*S*)-methyl 2-(tert-butoxycarbonylamino)-3-((*S*)-2-oxopyrrolidin-3-yl)propanoate (2).

In a hydrogenation flask was placed compound **1** (4.0 g, 12.7 mmol), 5 mL of chloroform and 60 mL of methanol before the addition of PtO_2_. The resulting mixture was stirred under hydrogen at 20°C for 12 h. Then the mixture was filtered over Celite to remove the catalyst. NaOAc (6.77 g, 25.5 mmol) was added to the filtrate before the resulting mixture was stirred at 60°C for 12 h. The reaction was quenched with water (30 mL). The suspension was extracted with ethyl acetate. The organic layers were combined, dried (MgSO_4_), and filtered. The light-brown filtrate was concentrated and purified by silica gel column chromatography (petroleum ether/ethyl acetate = 4/1) to give the product **2** (2.20 g, 61%) as white solid. ^1^H NMR (CDCl_3_) δ 6.02 (1H, br), 5.49 (1H, d, *J* = 7.8 Hz), 4.27-4.33 (1H, m), 3.72 (3H, s), 3.31-3.36 (2H, m), 2.40-2.49 (2H, m), 2.06-2.16 (1H, m), 1.77-1.89 (2H, m), 1.41 (9H, s). ESI-MS (*m*/*z*): 287 (M + H)^+^.

Synthesis of (*S*)-methyl 2-amino-3-((*S*)-2-oxopyrrolidin-3-yl)propanoate (3).

Compound **2** (1.0 g, 3.5 mmol) was dissolved in 10 mL dichloromethane (DCM), then 10 mL trifluoroacetic acid (TFA) was added. The reaction mixture was stirred at 20°C for 0.5 h, and concentrated *in vacuo* to get a colorless oil, which could be used for the following step without purification.

ESI-MS (*m*/*z*): 187 (M + H)^+^.

Synthesis of methyl N-substituted amino-acid esters 4

#### General procedure

The methyl amino-acid ester hydrochloride (6.0 mmol) was dissolved in 20 mL CH_2_Cl_2_, and then acyl chloride (6.0 mmol), triethylamine (1.69 mL, 12.0 mmol) were added, before the reaction was stirred for 2 h at 20°C. The reaction mixture was diluted with 20 mL CH_2_Cl_2_, washed with 50 mL of saturated brine (2 × 25 mL), and dried over Na_2_SO_4_. The solvent was evaporated and the product **4** was obtained as white solid (70-95% yield), which could be used for the next step without further purification.

(S)-methyl 2-cinnamamido-3-phenylpropanoate (4a).

The methyl *L*-phenylalaninate hydrochloride (1.30 g, 6.0 mmol) was dissolved in 20 mL CH_2_Cl_2_, and then cinnamoyl chloride (1.00 g, 6.0 mmol), triethylamine (1.69 mL, 12.0 mmol) were added, before the reaction was stirred for 2 h at room temperature. The reaction mixture was diluted with 20 mL CH_2_Cl_2_, washed with 50 mL of saturated brine (2×25 mL), and dried over Na_2_SO_4_. The solvent was evaporated and the product **4a** was obtained as white solid (1.75 g, 95%), which could be used for the next step without further purification.

#### Synthesis of N-substituted amino acids 5 (general procedure)

1 M NaOH (5 mL) was added to a solution of compound **4** (3.0 mmol) in methanol (5 mL). The reaction was stirred for 20 min at 20°C. Then 1 M HCl was added to the reaction solution until pH = 1. Then the reaction mixture was extracted with 100 mL of CH_2_Cl_2_ (2 × 50 mL) and the organic layer was washed with 50 mL of brine and dried over Na_2_SO_4_. The solvent was evaporated and the crude material purified on silica, eluted with mixtures of CH_2_Cl_2_/MeOH (20/1) to afford the product **5** (90-96% yield) as a white solid.

#### Synthesis of compounds 6 (General procedure)

Compound **5** (2.7 mmol) was dissolved in 10 mL of dry CH_2_Cl_2_. To this solution, 1.5 equiv (1.54 g) of 1-[bis(dimethylamino)methylene]-1H-1,2,3-triazolo[4,5-b]pyridinium 3-oxide hexafluorophosphate (HATU) was added, and the reaction was stirred for 0.5 h at 20°C. Then compound **3** (500 mg, 2.7 mmol) and TEA (0.70 mL, 5.42 mmol) was added to the reaction. The reaction was stirred for another 6 h. The reaction mixture was poured into 10 mL water. The aqueous solution was extracted with 50 mL of CH_2_Cl_2_ (2 × 25 mL) and washed with 50 mL of saturated brine (2 × 25 mL) and dried over Na_2_SO_4_. The solvent was evaporated and the crude material purified on silica, eluted with a mixture of CH_2_Cl_2_/MeOH (40/1) to give the product **6** (62-84% yield).

#### Synthesis of alcohols 7 (general procedure)

Compound **6** (1.1 mmol) was dissolved in methanol (40 mL), then NaBH_4_ (0.34 g, 8.8 mmol) was added under ambient conditions. The reaction mixture was stirred at 20°C for 2 h. Then the reaction was quenched with water (30 mL). The suspension was extracted with ethyl acetate. The organic layers were combined, dried, and filtered. The filtrate was evaporated to dryness and could be used for the next step without further purification (46-85% yield).

#### Synthesis of aldehydes 8 (general procedure)

Compound **7** (0.75 mmol) was dissolved in CH_2_Cl_2_, then Dess-Martin periodinane (337 mg, 0.79 mmol) and NaHCO_3_ (66 mg, 0.79 mmol) were added. The resulting mixture was stirred at 20°C for 1 h. The mixture was concentrated and purified by column chromatography on silica gel (CH_2_Cl_2_/MeOH = 20/1) to give the product **8** as white solid (88-95% yield).

#### Synthesis of compounds 9 (general procedure)

Compound **8** (0.40 mmol) was dissolved in CH_2_Cl_2_, and then acetic acid (0.028 g, 0.47 mmol) and isocyanide (0.43 mmol) were added successively to the solution. The reaction was stirred at 20°C for 24 h. Then the solvent was evaporated and the crude material purified on silica, eluted with a mixture of CH_2_Cl_2_/MeOH (20/1) to give the product **9** (46-84%).

#### Synthesis of α-hydroxyamides 10 (general procedure)

1 M NaOH (0.5 mL) was added to a solution of compound **9** (0.164 mmol) in methanol (5 mL). The reaction was stirred at 20°C for 0.5 h until the consumption of compound **9** was confirmed by TLC analysis. Then, 1 M HCl was added to the reaction solution until pH = 7. Following this, the solvent was evaporated to generate the product **10** as white solid, which could be used directly in the next step.

#### Synthesis of α-ketoamides 11 (general procedure)

Compound **10** was dissolved in CH_2_Cl_2_, then Dess-Martin periodinane (74 mg, 0.176 mmol) and NaHCO_3_ (30 mg, 0.176 mmol) were added. The resulting mixture was stirred at 20°C for 1 h. The mixture was concentrated and purified by column chromatography on silica gel (CH_2_Cl_2_/MeOH = 20/1) to give the α-ketoamides **11** as light yellow solid (52-79% in two steps).

(*S*)-*N*-benzyl-3-((*S*)-2-cinnamamido-3-phenylpropanamido)-2-oxo-4-((*S*)-2-oxopyrrolidin-3-yl)butanamide (11a).

75% yield. ^1^H NMR (400 MHz, CDCl_3_) δ 7.92 (d, *J* = 7.6 Hz, 1H), 7.56 (d, *J* = 15.2 Hz, 1H), 7.45-7.43 (m, 2H), 7.35-7.19 (m, 12H), 7.02-6.98 (m, 1H), 6.48 (d, *J* = 15.2 Hz, 1H), 6.44-6.42 (m, 1H), 5.01-4.92 (m, 2H), 4.46 (d, *J* = 8.4 Hz, 2H), 3.25-3.03 (m, 4H), 2.24-2.21 (m, 2H), 1.95-1.86 (m, 1H), 1.74-1.69 (m, 1H), 1.55-1.49 (m, 1H) ppm. ESI-MS (*m*/*z*): 567 [M + H]^+^.

For compounds **11b - 11e**, see Supplementary Information.

*tert*-Butyl ((*S*)-4-(benzylamino)-3,4-dioxo-1-((*S*)-2-oxopyrrolidin-3-yl)butan-2-yl)carbamate (11f)

^1^H NMR (300 MHz, CDCl_3_) δ 7.32-7.22 (m, 5H), 6.47 (br, 1H), 5.10 (d, *J* = 8.4 Hz, 1H), 4.37-4.26 (m, 3H), 3.37-3.32 (m, 2H), 2.53-2.47 (m, 2H), 2.05-1.98 (m, 1H), 1.85-1.79 (m, 1H), 1.62-1.56 (m, 1H), 1.44 (s, 9H) ppm. ESI-MS (*m*/*z*): 390 [M + H]^+^.

For compounds **11g - 11l**, see Supplementary Information.

(*S*)-*N*-benzyl-3-((*S*)-2-cinnamamido-3-(4-fluorophenyl)propanamido)-2-oxo-4-((*S*)-2-oxopyrrolidin-3-yl)butanamide (11m)

78% yield, ^1^H NMR (400 MHz, CDCl_3_) δ 7.91 (d, *J* = 7.6 Hz, 1H), 7.55 (d, *J* = 15.2 Hz, 1H), 7.43-7.40 (m, 2H), 7.35-7.11 (m, 11H), 7.01-6.98 (m, 1H), 6.46 (d, *J* = 15.2 Hz, 1H), 6.35-6.31 (m, 1H), 4.99-4.91 (m, 2H), 4.43 (d, *J* = 8.8 Hz, 2H), 3.27-3.12 (m, 3H), 3.05-2.99 (m, 1H), 2.24-2.21 (m, 2H), 2.03-1.96 (m, 1H), 1.72-1.54 (m, 2H) ppm. ESI-MS (*m*/*z*): 585 [M + H]^+^.

(*S*)-*N*-((*S*)-4-(benzylamino)-3,4-dioxo-1-((*S*)-2-oxopyrrolidin-3-yl)butan-2-yl)-2-cinnamamido-4-methylpentanamide (11n)

57% yield, ^1^H NMR (400 MHz, CDCl_3_) δ 8.01 (d, *J* = 7.6 Hz, 1H), 7.58 (d, *J* = 15.6 Hz, 1H), 7.43-7.35 (m, 5H), 7.33-7.14 (m, 4H), 7.02-6.98 (m, 1H), 6.48 (d, *J* = 15.6 Hz, 1H), 6.37-6.32 (m, 1H), 4.94-4.86 (m, 1H), 4.68-4.62 (m, 1H), 4.46 (d, *J* = 8.4 Hz, 2H), 3.25-3.11(m, 1H), 3.09-3.06 (m, 1H), 2.25-2.21 (m, 2H), 1.99-1.92 (m, 1H), 1.73-1.64 (m, 3H), 1.58-1.48 (m, 2H), 0.92 (d, *J* = 8.4 Hz, 3H), 0.88 (d, *J* =8.4 Hz, 3H) ppm. ESI-MS (*m*/*z*): 533 [M + H]^+^.

(*S*)-*N*-((*S*)-4-(benzylamino)-3,4-dioxo-1-((*S*)-2-oxopyrrolidin-3-yl)butan-2-yl)-2-cinnamamidohexanamide (11o)

76% yield, ^1^H NMR (400 MHz, CDCl_3_) δ 7.91 (d, *J* = 7.6 Hz, 1H), 7.55 (d, *J* = 15.2 Hz, 1H), 7.43-7.36 (m, 5H), 7.28-7.14 (m, 4H), 7.01-6.98 (m, 1H), 6.45 (d, *J* = 15.2 Hz, 1H), 6.37-6.32 (m, 1H), 4.98-4.91 (m, 1H), 4.73-4.67 (m, 1H), 4.48 (d, *J* = 8.0 Hz, 2H), 3.25-3.11(m, 1H), 3.09-3.03 (m, 1H), 2.25-2.21 (m, 2H), 1.92-1.86 (m, 1H), 1.73-1.64 (m, 3H), 1.56-1.52 (m, 1H), 1.36-1.25 (m, 4H), 0.93 (t, *J* = 8.4 Hz, 3H) ppm. ESI-MS (*m*/*z*): 533 [M + H]^+^.

(*S*)-*N*-((*S*)-4-(benzylamino)-3,4-dioxo-1-((*S*)-2-oxopyrrolidin-3-yl)butan-2-yl)-2-cinnamamidopent-4-ynamide (11p)

65% yield, ^1^H NMR (500 MHz, CDCl_3_) δ 8.56 (d, *J* = 7.5 Hz, 1H), 7.60 (d, *J* = 15.0 Hz, 1H), 7.53-7.46 (m, 2H), 7.38-7.17 (m, 7H), 6.53-6.42 (m, 2H), 5.32-5.25 (m, 1H), 4.85-4.65 (m, 1H), 4.47 (d, *J* = 8.5 Hz, 2H), 3.43-3.29 (m, 3H), 2.59-2.45 (m, 1H), 2.20-1.60 (m, 7H) ppm. ESI-MS (*m*/*z*): 515 [M + H]^+^.

(*S*)-*N*-benzyl-3-((*S*)-2-cinnamamido-2-cyclopropylacetamido)-2-oxo-4-((*S*)-2-oxopyrrolidin-3-yl)butanamide (11q).

66% yield, ^1^H NMR (500 MHz, CDCl_3_) δ 8.62 (d, *J* = 7.5 Hz, 1H), 7.60 (d, *J* = 15.0 Hz, 1H), 7.53-7.43 (m, 2H), 7.35-7.17 (m, 7H), 6.76-6.69 (m, 1H), 6.59-6.48 (m, 1H), 5.35-5.25 (m, 1H), 4.85-4.72 (m, 1H), 4.48 (d, *J* = 8.5 Hz, 2H), 3.38-3.22 (m, 2H), 2.62-2.45 (m, 1H), 2.12-1.63 (m, 4H), 1.20-0.92 (m, 1H), 0.46 (t, *J* = 7.0 Hz, 2H), 0.16-0.07 (m, 2H) ppm. ESI-MS (*m*/*z*): 517 [M + H]^+^.

(*S*)-*N*-benzyl-3-((*S*)-2-cinnamamido-3-cyclohexylpropanamido)-2-oxo-4-((*S*)-2-oxopyrrolidin-3-yl)butanamide (11r)

71% yield, ^1^H NMR (500 MHz, CDCl_3_) δ 8.56 (t, *J* = 6.0 Hz, 1H), 7.61 (d, *J* = 16.0 Hz, 1H), 7.52-7.44 (m, 3H), 7.35-7.20 (m, 6H), 6.66-6.59 (m, 1H), 6.48 (d, *J* = 13.0 Hz, 1H), 5.32-5.27 (m, 1H), 4.95-4.75 (m, 1H), 4.48 (d, *J* = 6.5 Hz, 2H), 3.39-3.29 (m, 2H), 2.65-2.35 (m, 2H), 2.09-1.68 (m, 10H), 1.29-1.16 (m, 4H), 1.00-0.88 (m, 2H) ppm. ESI-MS (*m*/*z*): 573 [M + H]^+^.

(*S*)-*N*-benzyl-3-((*S*)-2-cinnamamido-3-cyclopropylpropanamido)-2-oxo-4-((*S*)-2-oxopyrrolidin-3-yl)butanamide (11s).

64% yield, ^1^H NMR (500 MHz, CDCl_3_) δ 8.64 (d, *J* = 7.5 Hz, 1H), 7.60 (d, *J* = 15.0 Hz, 1H), 7.52-7.46 (m, 2H), 7.36-7.17 (m, 7H), 6.54-6.42 (m, 2H), 5.35-5.25 (m, 1H), 4.85-4.75 (m, 1H), 4.46 (d, *J* = 8.5 Hz, 2H), 3.38-3.29 (m, 2H), 2.65-2.35 (m, 1H), 2.15-1.90 (m, 2H), 1.85-1.60 (m, 4H), 0.90-0.72 (m, 1H), 0.47 (t, *J* = 7.0 Hz, 2H), 0.15-0.07 (m, 2H) ppm. ESI-MS (*m*/*z*): 531 [M + H]^+^.

(*S*)-*N*-benzyl-3-((*S*)-2-cinnamamido-3-cyclobutylpropanamido)-2-oxo-4-((*S*)-2-oxopyrrolidin-3-yl)butanamide (11t)

77% yield, ^1^H NMR (500 MHz, CDCl_3_) δ 8.62 (t, *J* = 6.5 Hz, 1H),7.60 (d, *J* = 15.0 Hz, 1H), 7.52-7.46 (m, 2H), 7.35-7.19 (m, 7H), 6.72-6.60 (m, 1H), 6.48 (d, *J* = 15.0 Hz, 1H), 5.32-5.26 (m, 1H), 4.77-4.69 (m, 1H), 4.49 (d, *J* = 6.5 Hz, 2H), 3.40-3.31 (m, 2H), 2.60-2.35 (m, 3H), 2.09-1.68 (m, 11H) ppm. ESI-MS (*m*/*z*): 545 [M + H]^+^.

(*S*)-*N*-benzyl-3-((*S*)-2-cinnamamido-3-cyclopentylpropanamido)-2-oxo-4-((*S*)-2-oxopyrrolidin-3-yl)butanamide (11u)

79% yield, ^1^H NMR (500 MHz, CDCl_3_) δ 7.60 (d, *J* = 15.0 Hz, 1H), 7.50-7.44 (m, 2H), 7.36-7.20 (m, 7H), 6.76-6.69 (m, 1H), 6.59-6.48 (m, 1H), 5.35-5.27 (m, 1H), 4.95-4.65 (m, 1H), 4.45 (d, *J* = 6.5 Hz, 2H), 3.38-3.29 (m, 2H), 2.65-2.35 (m, 1H), 2.00-1.38 (m, 13H), 1.20-1.00 (m, 2H) ppm. ESI-MS (*m*/*z*): 559 [M + H]^+^.

## ASSOCIATED CONTENT

### Supporting Information

Detailed results of the variation of the P1’ and P3 substituents

Synthesis of α-ketoamides **11b** - **11e** and **11g** - **11l** Supplemental Table 1: inhibitory activities (IC_50_ (μM)) of α-ketoamides with P1’ and P3 modifications against viral proteases

Supplemental Table 2: crystallographic data for complexes between viral proteases and α-ketoamides

Molecular formula strings and biological data (CSV)

### Accession Codes

Authors will release the atomic coordinates and experimental data upon article publication. Atomic coordinates include SARS-CoV M^pro^ in complex with compounds **11a** (5N19), **11s** (5N5O), HCoV-NL63 M^pro^ in complex with **11a** (6FV2), **11n** (6FV1), **11f** (5NH0), and CVB3 3C^pro^ in complex with (5NFS).

## Author Contributions

The manuscript was written through contributions by all authors. / All authors have given approval to the final version of the manuscript. / ‡These authors contributed equally.

## ACKNOWLEDGMENT

We thank Doris Mutschall, Javier Carbajo-Lozoya, Dev Raj Bairad, and Sebastian Schwinghammer for expert technical assistance, Dr. Bo Zhang and Prof. Frank van Kuppeveld (Wuhan and Utrecht, resp.) for the EV-A71 and CVB3 replicons, Prof. Ron Fouchier (Rotterdam) for the EMC/2012 strain of MERS-CoV, and Prof. Volker Thiel (Berne) for HCoV 229E. We are grateful to Dr. Naoki Sakai and the staff at synchrotron beamlines for help with diffraction data collection.

## ABBREVIATIONS

3CL^pro^: 3C-like protease;
3C^pro^: 3C protease;
A_490_: absorbance at 490 nm;
*ap*: antiperiplanar;
BAC: bacterial artificial chromosome;
CPE: cytopathic effect;
CVA16: Coxsackievirus A16;
CVB3: Coxsackievirus B3;
DMEM: Dulbecco’s modified minimal essential medium;
EMEM: Eagle’s minimal essential medium;
EV: enterovirus;
FIPV: Feline Infectious Peritonitis Virus;
FRET: fluorescence resonance energy transfer;
GlnLactam: glutamine lactam;
HCoV: human coronavirus;
HFMD: Hand, Foot, and Mouth Disease;
HRV: human rhinovirus;
MERS-CoV: Middle-East respiratory syndrome coronavirus;
M^pro^: main protease;
Nsp5: non-structural protein 5;
NTR: non-translated region;
OD_498_: optical density at 498 nm;
PEG: polyethylene glycol;
RFU: relative fluorescence units;
Rz: ribozyme;
SARS: severe acute respiratory syndrome;
SARS-CoV: SARS coronavirus;
- *sc*: (-)-synclinal;
SD: standard deviation;
TLC: thin-layer chromatography

## SYNOPSIS TOC

### Table of Contents Graphic

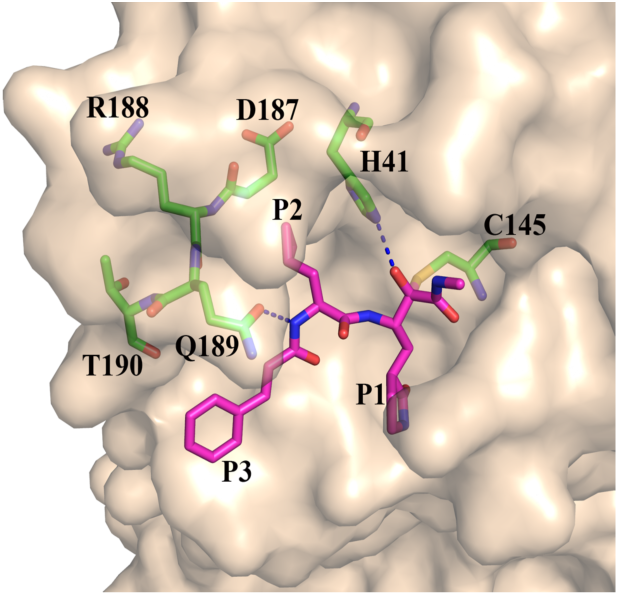

