## SUPPORTING INFORMATION for "Alpha-ketoamides as broad-spectrum inhibitors of coronavirus and enterovirus replication"

*Structure-based design, synthesis, and activity assessment*

Linlin Zhang^1,2,‡^, Daizong Lin^1,2,3,‡,†^, Yuri Kusov^1^, Yong Nian^3^, Qingjun Ma^1^, Jiang Wang^3^, Albrecht von Brunn^4^, Pieter Leyssen^5^, Kristina Lanko^5^, Johan Neyts^5^, Adriaan de Wilde^6^, Eric J. Snijder^6^, Hong Liu^3^*, Rolf Hilgenfeld^1,2,3^*

^1^Institute of Biochemistry, Center for Structural and Cell Biology in Medicine, University of Lübeck, 23562 Lübeck, Germany.

^2^German Center for Infection Research (DZIF), Hamburg - Lübeck - Borstel - Riems Site, University of Lübeck, Lübeck, Germany.

^3^Shanghai Institute of Materia Medica, 201203 Shanghai, China

^4^Max von Pettenkofer Institute, Ludwig-Maximilians-University Munich, 80336 Munich, Germany

^5^Rega Institute for Medical Research, University of Leuven, 3000 Leuven, Belgium

^6^Leiden University Medical Center, 2333 ZA Leiden, The Netherlands

^‡^These authors contributed equally.

**Table of Contents**

**1. Supplementary Results	2**

**1.1 Variation of the P1' and P3 substituents	2**

**1.1.1 P1' substituent	2**

**1.1.2 P3 substituent	2**

**2. Supplementary Experimental Section	2**

**2.1 Synthesis of α-ketoamides 11b - 11e and 11g - 11l	2**

**3. Supplementary Tables	4**

**3.1 Supplementary Table 1	4**

**3.1 Supplementary Table 2	6**

**SUPPLEMENTARY RESULTS**

**Variation of the P1' and P3 substituents**

***P1' substituent***

The crystal structures indicated that the fit of the P1' benzyl group of **11a** in the S1' pocket might be improved. The amide oxygen accepts an H-bond from the imidazole of His41 in the complexes with SARS-CoV M^pro^ and HCoV-NL63 M^pro^, and from the oxyanion-hole amide of Gly145 in CVB3 3C^pro^, but the P1' benzyl group appears not fully embedded in the pocket (see Fig. 3a-c). We therefore replaced it by n-butyl in **11b** (Table S1). This resulted in an improvement of the inhibitory activity against CVB3 3C^pro^ (from IC_50_ = 6.6 to 1.0 μM), whereas that against the EV-A71 enzyme was somewhat weaker (IC_50_ = 2.4 μM). Most importantly, however, the compound proved totally inactive against recombinant SARS-CoV M^pro^ (IC_50_ > 50 μM). Replacement of the P1' substituent by *tert*-butyl in **11c** led to very poor activities against all three proteases, and P1' = isobutyl (**11d**) or 2-methoxy-2-oxoethyl (**11e**) were not much better. Therefore, although there is probably room for further improvement, we decided to maintain the original design with P1' = benzyl.

***P3 substituent***

Inspection of the crystal structures of **11a** in complex with SARS-CoV M^pro^ and CVB3 3C^pro^ suggested that the interaction of the P3-cinnamoyl group with the binding cleft might be sub-optimal (Fig. 3a,c). In order to furnish the P3 group with more flexibility, we reduced the double bond in **11a** to generate **11g**. However, presumably for entropic reasons, this led to a 2- to 3-fold decrease in activity against the SARS-CoV M^pro^ and the EV-A71 3C^pro^, compared to compound **11a** (Table S1). CVB3 3C^pro^ was the only protease against which an improvement by the reduction of the double bond of the cinnamoyl group was noted, but in the replicon assays, compound **11g** was inferior to **11a** in all cases (Table S2).

Introduction in P3 of less flexible, bulky substituents such as 2,3-dihydrobenzo[b][1,4]dioxine-6-carbonyl (**11h**) and 2,3-dihydrobenzo[b][1,4]dioxine-5-carbonyl (**11i**) also met with limited success; IC_50_ values were mediocre for inhibition of the EV-A71 3C^pro^ and the CVB3 3C^pro^ and higher by a factor of at least 5 for inhibition of the SARS-CoV M^pro^, compared to compound **11a**. P3 = benzofuran-2-carbonyl (**11j**) led to a further reduction of inhibitory activity, particularly against the EV-A71 3C^pro^. Compounds with P3 = benzo[b]thiophene-2-carbonyl (**11k**) and 6-bromoimidazo[1,2-a]pyridine-2-carbonyl (**11l**) were inactive against EV-A71 3C^pro^ and only very moderately active against CVB3 3C^pro^ and SARS-CoV M^pro^ (Table S1). Inspection of the template crystal structures revealed that all of these substituents would occupy the space of the main chain of a peptide substrate at the S3 and partly S4 sites and are unable to reach into the space normally taken by a P3 amino-acid side-chain (which would be relatively unrestricted in all three proteases). However, in the predicted main-chain-like orientation, the extended aromatic groups would interfere with the NH group of Leu127 and the carbonyl of Gly164.

In summary, the cinnamoyl group of the original compound, **11a**, was superior to all other substituents explored. Among the compounds tested thus far, **11a** resulted in the best inhibitory activities towards both enterovirus 3C^pro^ and SARS-CoV M^pro^. Accordingly, we decided to retain the cinnamoyl group as the substituent at the P3 position.

**SUPPLEMENTARY EXPERIMENTAL SECTION**

***Synthesis of a-ketoamides 11b - 11e and 11g - 11l***

**(*S*)-*N*-butyl-3-((*S*)-2-cinnamamido-3-phenylpropanamido)-2-oxo-4-((*S*)-2-oxopyrrolidin-3-yl)butanamide (11b)**

62% yield, ^1^H NMR (400 MHz, CDCl_3_) δ 7.78 (d, *J* = 7.6 Hz, 1H), 7.56 (d, *J* = 16.0 Hz, 1H), 7.46-7.43 (m, 2H), 7.32-7.30 (m, 3H), 7.28-7.14 (m, 4H), 7.01-6.98 (m, 1H), 6.53 (d, *J* = 16.0 Hz, 1H), 6.42-6.40 (m, 1H), 5.01-4.92 (m, 2H), 3.44-3.03 (m, 6H), 2.47-2.37 (m, 2H), 1.95-1.84 (m, 1H), 1.74-1.71 (m, 1H), 1.55-1.51 (m, 3H), 1.32-1.28 (m, 2H), 0.90 (t, *J* = 8.0 Hz, 1H) ppm. ESI-MS (*m*/*z*): 533 [M + H]^+^.

**(*S*)-*N*-(*tert*-butyl)-3-((*S*)-2-cinnamamido-3-phenylpropanamido)-2-oxo-4-((*S*)-2-oxopyrrolidin-3-yl)butanamide (11c)**

55% yield, ^1^H NMR (400 MHz, CDCl_3_) δ 7.79 (d, *J* = 7.6 Hz, 1H), 7.57 (d, *J* = 16.0 Hz, 1H), 7.46-7.43 (m, 2H), 7.32-7.13 (m, 7H), 7.01-6.98 (m, 1H), 6.59 (d, *J* = 16.0 Hz, 1H), 5.02-4.92 (m, 2H), 3.24-3.03 (m, 4H), 2.47-2.37 (m, 2H), 1.95-1.84 (m, 1H), 1.74-1.71 (m, 1H), 1.55-1.51 (m, 1H), 1.33 (s, 9H) ppm. ESI-MS (*m*/*z*): 533 [M + H]^+^.

**(*S*)-*N*-((*S*)-4-(isobutylamino)-3,4-dioxo-1-((*S*)-2-oxopyrrolidin-3-yl)butan-2-yl)-2-cinnamamido-4-methylpentanamide (11d)**

72% yield, ^1^H NMR (400 MHz, CDCl_3_) δ 8.00 (d, *J* = 7.6 Hz, 1H), 7.54 (d, *J* = 15.6 Hz, 1H), 7.42-7.35 (m, 3H), 7.23 (d, *J* = 7.2 Hz, 2H), 7.00-6.96 (m, 1H), 6.48 (d, *J* = 15.6 Hz, 1H), 6.36-6.31 (m, 1H), 4.95-4.84 (m, 1H), 4.69-4.62 (m, 1H), 3.23-3.13 (m, 1H), 3.04-2.86 (m, 3H), 2.25-2.21 (m, 2H), 2.11-2.07 (m, 1H), 1.96-1.92 (m, 1H), 1.72-1.64 (m, 3H), 1.56-1.48 (m, 2H), 0.95-0.91 (m, 6H), 0.90-0.86 (m, 6H) ppm. ESI-MS (*m*/*z*): 499 [M + H]^+^.

**Methyl 2-((*S*)-3-((*S*)-2-cinnamamido-3-phenylpropanamido)-2-oxo-4-((*S*)-2-oxopyrrolidin-3-yl)butanamido)acetate (11e)**

52% yield, ^1^H NMR (400 MHz, CDCl_3_) δ 7.78 (d, *J* = 7.6 Hz, 1H), 7.56 (d, *J* = 16.0 Hz, 1H), 7.48-7.42 (m, 2H), 7.37-7.13 (m, 7H), 7.01-6.99 (m, 1H), 6.53 (d, *J* = 16.0 Hz, 1H), 6.42-6.40 (m, 1H), 5.03-4.95 (m, 2H), 4.42-4.14 (m, 4H), 3.68 (s, 3H), 3.24-3.03 (m, 4H), 2.42-2.34 (m, 2H), 1.95-1.83 (m, 1H), 1.76-1.71 (m, 1H), 1.56-1.52 (m, 1H) ppm. ESI-MS (*m*/*z*): 549 [M + H]^+^.

**(*S*)-*N*-benzyl-2-oxo-4-((*S*)-2-oxopyrrolidin-3-yl)-3-((*S*)-3-phenyl-2-(3-phenylpropanamido)propanamido)butanamide (11g)**

73% yield, ^1^H NMR (400 MHz, CDCl_3_) δ 7.92 (d, *J* = 7.6 Hz, 1H), 7.46-7.43 (m, 2H), 7.35-7.13 (m, 12H), 7.01-6.96 (m, 1H), 6.45-6.42 (m, 1H), 4.97-4.91 (m, 2H), 4.46 (d, *J* = 8.4 Hz, 2H), 3.25-3.03 (m, 4H), 2.86-2.78 (m, 2H), 2.55 (t, *J* = 8.0 Hz, 2H), 2.26-2.22 (m, 2H), 1.95-1.86 (m, 1H), 1.74-1.69 (m, 1H), 1.55-1.48 (m, 1H) ppm. ESI-MS (*m*/*z*): 569 [M + H]^+^.

***N*-((*S*)-1-(((*S*)-4-(benzylamino)-3,4-dioxo-1-((*S*)-2-oxopyrrolidin-3-yl)butan-2-yl)amino)-1-oxo-3-phenylpropan-2-yl)-2,3-dihydrobenzo[*b*][1,4]dioxine-6-carboxamide (11h)**

64% yield, ^1^H NMR (400 MHz, CDCl_3_) δ 8.51 (d, *J* = 26.7 Hz, 1H), 7.59 – 7.27 (m, 4H), 7.26 – 6.95 (m, 11H), 6.80 – 6.71 (m, 1H), 5.06 (d, *J* = 31.7 Hz, 1H), 4.41 (s, 2H), 4.18 (d, *J* = 8.6 Hz, 4H), 3.32 – 2.95 (m, 5H), 2.38 (d, *J* = 73.2 Hz, 1H), 2.21 – 1.69 (m, 4H). HRMS (ESI) *m*/*z*: calcd for C_33_H_35_N_4_O_7_ ^+^ [M + H]^+^ : 599.2506, found: 599.2501.

***N*-((*S*)-1-(((*S*)-4-(benzylamino)-3,4-dioxo-1-((*S*)-2-oxopyrrolidin-3-yl)butan-2-yl)amino)-1-oxo-3-phenylpropan-2-yl)-2,3-dihydrobenzo[*b*][1,4]dioxine-5-carboxamide (11i)**

60% yield, ^1^H NMR (400 MHz, CDCl_3_) δ 8.17 – 8.02 (m, 2H), 7.67 – 7.60 (m, 1H), 7.43 – 7.26 (m, 6H), 7.26 – 7.16 (m, 5H), 6.99 – 6.93 (m, 1H), 6.87 (tt, *J* = 4.9, 2.4 Hz, 1H), 6.05 (dd, *J* = 52.8, 27.5 Hz, 1H), 5.40 – 5.28 (m, 1H), 5.09 – 4.97 (m, 1H), 4.52 – 4.41 (m, 2H), 4.30 – 4.17 (m, 4H), 3.34 – 3.24 (m, 2H), 3.16 (ddd, *J* = 13.8, 8.1, 3.6 Hz, 2H), 2.51 – 2.40 (m, 1H), 2.28 – 2.12 (m, 1H), 2.08 – 1.94 (m, 2H), 1.86 (dd, *J* = 12.8, 6.1 Hz, 1H). HRMS (ESI) *m*/*z*: calcd for C_33_H_35_N_4_O_7_ ^+^ [M + H]^+^ : 599.2506, found: 599.2499.

***N*-((*S*)-1-(((*S*)-4-(benzylamino)-3,4-dioxo-1-((*S*)-2-oxopyrrolidin-3-yl)butan-2-yl)amino)-1-oxo-3-phenylpropan-2-yl)benzofuran-2-carboxamide (11j)**

72% yield, ^1^H NMR (400 MHz, CDCl_3_) δ 8.75 – 8.54 (m, 1H), 7.70 – 7.55 (m, 2H), 7.52 – 7.38 (m, 4H), 7.37 – 7.26 (m, 7H), 7.25 – 7.14 (m, 3H), 6.66 (dt, *J* = 20.2, 8.9 Hz, 1H), 5.49 – 5.29 (m, 1H), 5.23 – 5.07 (m, 1H), 4.58 – 4.41 (m, 2H), 3.36 – 3.10 (m, 4H), 2.57 – 2.39 (m, 1H), 2.31 – 2.21 (m, 1H), 2.14 – 1.83 (m, 3H). HRMS (ESI) *m*/*z*: calcd for C_33_H_33_N_4_O_6_ ^+^ [M + H]^+^ : 581.2400, found: 581.2388.

***N*-((*S*)-1-(((*S*)-4-(benzylamino)-3,4-dioxo-1-((*S*)-2-oxopyrrolidin-3-yl)butan-2-yl)amino)-1-oxo-3-phenylpropan-2-yl)benzo[*b*]thiophene-2-carboxamide** **(11k)**

66% yield, ^1^H NMR (400 MHz, CDCl_3_) δ 8.75 – 8.42 (m, 1H), 7.77 (tdd, *J* = 17.9, 11.8, 5.7 Hz, 3H), 7.44 – 7.26 (m, 8H), 7.20 (ddd, *J* = 16.2, 10.1, 5.6 Hz, 5H), 7.12 – 6.92 (m, 1H), 6.26 – 5.93 (m, 1H), 5.46 – 5.21 (m, 1H), 5.16 – 4.92 (m, 1H), 4.52 – 4.39 (m, 2H), 3.35 – 3.07 (m, 5H), 2.50 – 2.34 (m, 1H), 2.26 – 2.17 (m, 1H), 2.13 – 1.85 (m, 3H). HRMS (ESI) *m*/*z*: calcd for C_33_H_33_N_4_O_5_S ^+^ [M + H]^+^ : 597.2172, found: 597.2179.

***N*-((*S*)-1-(((*S*)-4-(benzylamino)-3,4-dioxo-1-((*S*)-2-oxopyrrolidin-3-yl)butan-2-yl)amino)-1-oxo-3-phenylpropan-2-yl)-6-bromoimidazo[1,2-*a*]pyridine-2-carboxamide (11l)**

73% yield, ^1^H NMR (400 MHz, CDCl_3_) δ 8.38 – 7.82 (m, 5H), 7.46 – 7.27 (m, 5H), 7.25 – 7.07 (m, 7H), 6.58 – 6.21 (m, 1H), 5.04 (dd, *J* = 46.7, 39.6 Hz, 2H), 4.49 – 4.37 (m, 2H), 3.31 – 3.11 (m, 3H), 2.39 (d, *J* = 9.2 Hz, 1H), 2.25 – 2.16 (m, 2H), 2.02 – 1.97 (m, 1H), 1.87 (s, 1H). HRMS (ESI) *m*/*z*: calcd for C_32_H_32_BrN_6_O_5_ ^+^ [M + H]^+^ : 659.1618, found: 659.1624.

SUPPLEMENTARY TABLES

**Supplementary Table 1: Inhibitory activities (IC_50_ (μM)) of α-ketoamides with P1' and P3 modifications against viral proteases**

| **Compound No.** | **Formula** | **EV-A71 3C^pro^** | **CVB3 3C^pro^** | **SARS-CoV M^pro^** | **HCoV-NL63 M^pro^** |
| --- | --- | --- | --- | --- | --- |
| **11b** | 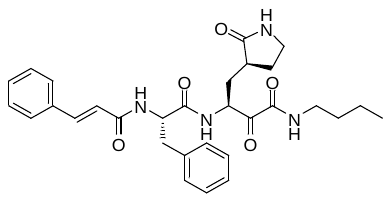 | 2.38 ± 0.58 | 1.00 ± 0.35 | >50 | nd |
| **11c** | 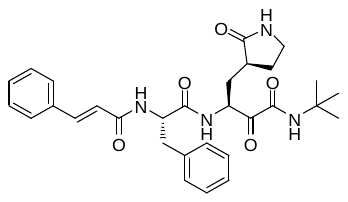 | 19.04 ± 5.79 | 9.86 ± 2.68 | 43.19 ± 24.68 | nd |
| **11d** | 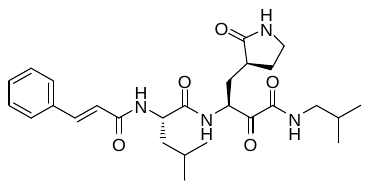 | 17.66 ± 8.78 | 6.50 ± 0.58 | 11.37 ± 4.64 | nd |
| **11e** | 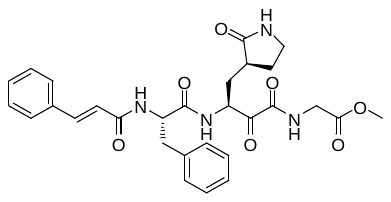 | 11.14 ± 2.64 | 9.91 ± 1.84 | 4.47 ± 2.49 | nd |
| **11g** | 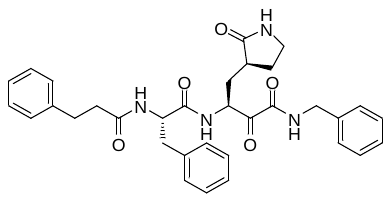 | 4.57 ± 1.04 | 3.50 ± 1.06 | 3.59 ± 0.24 | nd |
| **11h** | 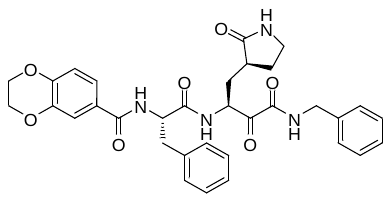 | 4.75 ± 2.73 | 3.66 ± 0.79 | 10.73 ± 3.55 | nd |
| **11i** | 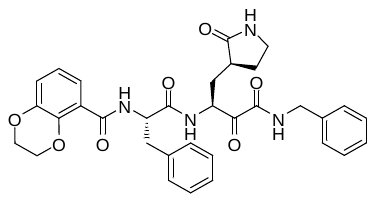 | 9.74 ± 3.88 | 4.82 ± 1.17 | 11.88 ± 5.29 | nd |
| **11j** | 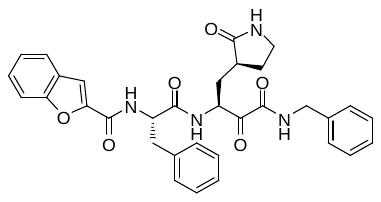 | 11.30 ± 6.52 | 9.19 ± 2.99 | 16.71 ± 6.96 | nd |
| **11k** | 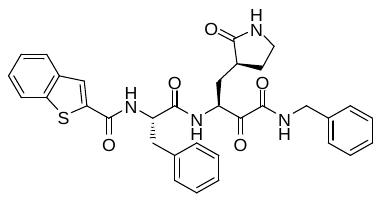 | >50 | 18.56 ± 0.58 | 9.68 ± 2.91 | nd |
| **11l** | 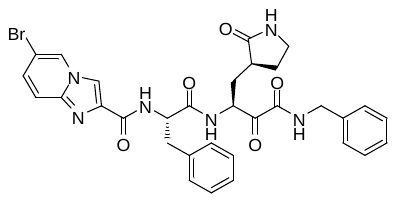 | >50 | 30.78 ± 12.92 | 13.25 ± 1.09 | nd |
| nd-not done |  |  |  |  |  |

**Supplementary Table 2: Crystallographic data for complexes between viral proteases and α-ketoamides**

| **Protease** | **SARS-CoV M^pro^** | **SARS-CoV M^pro^** | **HCoV-NL63 M^pro^** | **HCoV-NL63 M^pro^** | **HCoV-NL63 M^pro^** | **CVB3 3C^pro^** |
| --- | --- | --- | --- | --- | --- | --- |
| **Ligand** | **11a** | **11s** | **11a** | **11n** | **11f** | **11a** |
| **PDB entry** | 5N19 | 5N5O | 6FV2 | 6FV1 | 5NH0 | 5NFS |
| **Data collection statistics** |  |  |  |  |  |  |
| X-ray source | SLS X10SA | DESY P11 | BESSY 14.2 | BESSY 14.2 | BESSY 14.2 | BESSY 14.2 |
| Wavelength [Å] | 1.0000 | 0.9919 | 0.9184 | 0.9184 | 0.9184 | 0.9184 |
| Space group | *C*2 | *C*2 | *C*222_1_ | *C*222_1_ | *C*222_1_ | *C*2 |
| Unit cell dimensions [Å] | *a* = 108.20,  *b* = 82.28,  *c* = 53.44 | *a* = 109.17,  *b* = 81.87,  *c* = 53.50 | *a* = 131.12,  *b* = 211.04,  *c* = 115.63 | *a* = 133.86,  *b* = 211.33,  *c* = 118.32 | *a* = 133.65,  *b* = 211.29,  *c* = 118.26 | *a* = 77.66,  *b* = 64.35,  *c* = 39.48 |
| Unit cell angles [º] | α = γ = 90,  β = 103.95 | α = γ = 90,  β = 104.34 | α = β = γ = 90 | α = β = γ = 90 | α = β = γ = 90 | α = γ = 90  β = 115.44 |
| Number of protein molecules per asymmetric unit | 1 | 1 | 3 | 3 | 3 | 1 |
| Resolution range ^a^ [Å] | 64.77 - 1.62  (1.71 - 1.62) | 43.83 – 2.00  (2.11 - 2.00) | 48.00 - 2.95  (3.11 – 2.95) | 48.24 – 2.30  (2.42 -2.30) | 44.28 – 2.35  (2.48 - 2.35) | 33.56 – 1.80  (1.90 -1.80) |
| Number of observations | 386,723 (56,621) | 98,989 (14,526) | 188,558 (28,512) | 372,476 (54,396) | 301,104 (37,670) | 50,844 (7,025) |
| Number of unique reflections | 56,530 (8,106) | 30,329 (4,358) | 33,470 (4,925) | 74,507 (10,793) | 68,423 (9,796) | 16,209 (2,332) |
| Completeness [％] | 98.2 (97.4) | 98.3 (97.4) | 98.1 (99.8) | 99.9 (100.0) | 98.2 (97.2) | 99.4 (99.2) |
| Mean I/σ(I) | 24.5 (3.4) | 10.5 (2.4) | 13.1 (3.0) | 20.4 (3.3) | 11.5 (3.0) | 18.9 (2.9) |
| Multiplicity | 6.8 (7.0) | 3.3 (3.3) | 5.6 (5.8) | 5.0 (5.0) | 4.4 (3.8) | 3.1 (3.0) |
| R_merge_ ^b^ [%] | 0.034 (0.490) | 0.060 (0.507) | 0.112 (0.569) | 0.060 (0.536) | 0.076 (0.458) | 0.034 (0.393) |
| CC_1/2_ | 1.000 (0.960) | 0.997 (0.804) | 0.994 (0.808) | 0.999 (0.896) | 0.996 (0.873) | 0.999 (0.887) |
| **Protease** | **SARS-CoV M^pro^** | **SARS-CoV M^pro^** | **HCoV-NL63 M^pro^** | **HCoV-NL63 M^pro^** | **HCoV-NL63 M^pro^** | **CVB3 3C^pro^** |
| **Ligand** | **11a** | **11s** | **11a** | **11n** | **11f** | **11a** |
| **Refinement statistics** |  |  |  |  |  |  |
| R_cryst_ ^d^ /R_free_ ^e^ [%] | 17.25/20.04 | 19.22/25.15 | 17.83/23.97 | 19.27/22.92 | 19.08/23.28 | 17.75/23.01 |
| r.m.s.d. in bond lengths (Å) | 0.0280 | 0.0182 | 0.0140 | 0.0187 | 0.0184 | 0.0226 |
| r.m.s.d. in bond angles (°) | 2.484 | 1.908 | 1.908 | 1.974 | 2.000 | 2.386 |
| Average B-factor for protein atoms (Å^2^) | 33.21 | 44.06 | 56.20 | 51.86 | 44.37 | 32.14 |
| Average B-factor for ligand atoms (Å^2^) | 45.08 | 52.11 | 64.69 | 48.14 | 43.98 | 77.56 |
| Average B-factor for water molecules (Å^2^) | 46.90 | 52.40 | 94.16 | 44.57 | 66.50 | 52.17 |
| Number of protein atoms | 2392 | 2378 | 6824 | 6807 | 6819 | 1404 |
| Number of ligand atoms | 42 | 39 | 126 | 117 | 84 | 42 |
| Number of water molecules | 415 | 215 | 239 | 474 | 472 | 86 |
| **Ramachandran plot** |  |  |  |  |  |  |
| Preferred regions (%) | 97.99 | 97.01 | 95.98 | 97.20 | 97.73 | 94.38 |
| Allowed regions (%) | 1.68 | 2.66 | 3.80 | 2.58 | 2.27 | 5.06 |
| Outlier regions (%) | 0.34 | 0.33 | 0.22 | 0.22 | 0.00 | 0.56 |

a The highest resolution shell is shown in parantheses.

b ${R_{\mathrm{merge}}=\sum_{hkl} \sum_{i=1}^{n} \left| I_{i}(hkl)-\bar{I}(hkl) \right|}/{\sum_{hkl} \sum_{i=1}^{n} I_{i}(hkl)}$

c ${R_{\mathrm{pim}}=\sum_{hkl} \sqrt{1/(n-1)}\sum_{i=1}^{n} \left| I_{i}\left( hkl \right)-\bar{I}\left( hkl \right) \right|}/{\sum_{hkl} \sum_{i=1}^{n} I_{i}\left( hkl \right)}$ (Ref. 66)

d${R_{\mathrm{cryst}}=\sum_{hkl} \left| F_{o}(hkl)-F_{c}(hkl) \right|}/{\sum_{hkl} \left| F_{o}\left( hkl \right) \right|}$

e R_free_ was calculated for a test set of reflections (5%) omitted from the refinement.
